# The Bacteria of a Fig Microcommunity

**DOI:** 10.1101/2024.11.22.624729

**Authors:** Gavin C. Woodruff, Kimberly A. Moser, John Wang

**Affiliations:** School of Biological Sciences, University of Oklahoma; Biodiversity Research Center, Academia Sinica

**Keywords:** Figs, fig wasps, microbial ecology, *Caenorhabditis*, *C*. *elegans*

## Abstract

Understanding the biotic drivers of diversity is a major goal of microbial ecology. One approach towards tackling this issue is to interrogate relatively simple communities that are easy to observe and perturb. Figs (syconia) of the genus *Ficus* represent such a system. Here, we describe the microbial communities of *Ficus septica* figs, which are associated with the nematode *Caenorhabditis inopinata* (the sister species of the *C. elegans* genetic model system). In 2019, 38 *Ficus septica* figs (across 12 plants in Taiwan) were dissected, and metadata such as foundress wasp number and nematode occupancy were collected for each fig. Suspensions derived from interior fig material and fig surface washes were prepared for 16S microbial metabarcoding. Over 3,000 OTUs were detected, and microbial communities were dominated by members of *Proteobacteria*, *Bacteroidota*, and *Actinobacteriota*. Although microbial communities of fig exteriors and interiors can be distinguished, levels of microbial alpha diversity were comparable across these areas of the fig. Nematodes likewise had no detectable impact on microbial alpha diversity, although nematodes were associated with a modest change in microbial community composition. A handful of OTUs (associated with the genera *Kosokonia*, *Ochobactrum*, and *Stenotrophomonas*) revealed potential differential abundance among figs varying in nematode occupancy. Additionally, foundress wasp number was negatively correlated with microbial alpha diversity. These findings set the stage for future studies that directly test the role of nematode and wasp occupancy on microbial communities, as well as investigations that probe nematode-microbe interactions through laboratory experiments. Taken together, these results constitute a fundamental step in characterizing the natural microbial communities of figs and *Caenorhabditis* nematodes.

**Importance:** Unraveling why different species live in different places is a longstanding open question in ecology, and it is clear that interspecific interactions among species are a major contributor to species distributions. *Ficus* figs are a useful system for ecological studies because they are relatively simple microcosms where characterizing animal community composition of multiple samples is straightforward. Additionally, *Caenorhabditis inopinata*, a close relative of the *C. elegans* genetic model system, thrives in *Ficus septica* figs. Here, we tie 16S microbial metabarcoding to nematode and wasp occupancy data to understand the causes of bacterial community composition in *F. septica* figs. We found that microbial composition, but not total diversity, varies among fig surface and interiors. Likewise, we found that nematode occupancy impacts microbial composition but not alpha diversity. Moreover, we show that as the number of foundress wasps increases, the microbial alpha diversity decreases. Finally, we identified OTUs that are potentially associated with nematode occupancy. Taken together, these results represent a key step in describing a microcommunity wherein ecological genetic hypotheses can be tested, as well as one that can potentially reveal the roles of uncharacterized genes in established model systems.

## Introduction

Ecology seeks to explain species distributions. Biotic interactions are critical drivers of such distributions, as predator/prey, host/parasite, and interspecific competitive relationships (among others) contribute to the ability of an organism to thrive in a given place. However, while demonstrating that a given interaction causes species occupancy is notoriously difficult (e.g., demonstrating a host/parasite interaction has a large effect on a species distribution), even demonstrating the existence of an interspecific interaction in itself can be a challenge (Hortal et al. 2015; Morales-Castilla et al. 2015). Furthermore, simply noting the presence of an interaction is insufficient— the frequency and effect size of such interactions must be estimated to understand if they are relevant to species occupancy patterns. Moreover, manipulative experiments are often required to demonstrate causation, and manipulatable ecological systems are usually expensive (Hanson and Walker 2020). Thus, one avenue towards tackling ecological questions is through the use of systems where species interactions are easy to assess, quantify and manipulate, allowing empirical experiments.

Figs (also known as syconia) of the genus *Ficus* represent such a system. Although known in the common parlance as fruit (and mature figs are fruits indeed (Borges 2021)), figs are more appropriately described as an inflorescence with the flowers facing inward (Su 2008). Figs are pollinated by fig wasps, and fig wasps in turn lay their eggs in fig ovules (Borges 2021). The flowers are accessible to pollinators only via a specialized opening called the ostiole (Castro-Cárdenas et al. 2022). The ostiole is protected by a series of modified leaves (i.e., bracts) that presumably hinder animal penetration (Verkerke 1989; Castro-Cárdenas et al. 2022); these bracts can apparently cause the loss of antennae, wings, and appendages of pollinating wasp foundresses (Borges 2021; Hatta et al. 2023). While an extensive literature regarding the fig/fig wasp mutualism exists (Janzen 1979; Weiblen 2002; Segar et al. 2014; Borges 2015; Van Goor et al. 2023), more species interact with figs than pollinating wasps alone (although multiple species of fig wasp have been reported to pollinate a single fig (Su et al. 2008)). Non-pollinating fig wasps are common, with more than one species of wasp frequently parasitizing a single fig species (Borges 2015). Moreover, non-pollinating wasps can fill a variety of ecological niches within the fig microcommunity, living as gallers, wasp parasitoids, kleptoparasites, and seed eaters (Borges 2015). Various species of fungi have been observed in figs (Martinson et al. 2012). Ants are also associated with figs and can exert protective effects (Jandér 2015). Likewise, moth larvae can prey on fig tissue, potentially impacting community members (Sugiura and Yamazaki 2004). Additionally, mites (Jauharlina et al. 2012), nematodes (Van Goor et al. 2023), single celled eukaryotes, and bacteria also contribute to the fig microcommunity (Dong et al. 2022; Li et al. 2022).

Additionally, this complex community is also amenable to ecological work. Figs are small, discrete, individualized communities where species occupancies and abundances can be easily ascertained. Additionally, multiple figs from a single individual plant can be easily sampled and observed, affording reproducibility and large sample sizes. And, fitness proxies for multiple community members can be assessed (seed count, wasp progeny number, etc.) (Borges 2021). While the community is complex, the species composition of figs is not so extreme as to make the number of possible interactions intractable— that is, it strikes a balance between system complexity and observational feasibility. Furthermore, this system can be experimentally manipulated in various ways. For instance, nets can be placed over figs to prevent wasp entry (Liu et al. 2013). Alternatively, antibiotic, antifungal, or anthelmintic compounds could plausibly be added to figs to manipulate bacterial, fungal, or nematode occupancy. Additionally, as wasp (Borges 2015) and nematode (Susoy et al. 2016; Woodruff and Phillips 2018) demographics and occupancy can change across fig development, theories of ecological succession (Chang and Turner 2019) can also be applied to and tested with the fig system. For these reasons, figs are well-suited for ecological studies.

Nematodes are a particularly attractive subject of fig biology because of the widespread ecological diversity of fig nematodes (Van Goor et al. 2023). Nematode diversification has been extensive even within a single fig species (eight species of nematodes were observed within *F. burkei* (Kruger et al. 2021)). Nematode fig parasites (e.g., *Schistonchus*, *Martinema*, and *Ficophagus* (Davies et al. 2015)), wasp parasites (*Parasitodiplogaster* and *Teratodiplogaster* (POINAR and HERRE 1991; Kanzaki et al. 2009)), fungal feeders (*Bursaphelenchus* (Kanzaki et al. 2014)), bacterivores (*Caenorhabditis*, *Pristionchus*, and *Acrostichus* (Susoy et al. 2016; Kanzaki et al. 2018; Kruger et al. 2021)), and nematode predators (*Pristionchus*) all thrive in the fig microcosm. Additionally, all are thought to disperse on fig wasps to enable propagation in fresh figs. Consistent with this extensive ecological diversity, nematodes have evolved to live in figs at least eight times independently (Van Goor et al. 2023). The nematode *Caenorhabditis inopinata*, associated with *F. septica*, is of particular interest because it is the closest known relative of the widely-used genetic model system, *C. elegans* (Kanzaki et al. 2018).

Despite its fifty-year status as a genetic model (Brenner 1974), hundreds of *C. elegans* genes remain uncharacterized (Petersen et al. 2015). Presumably, one reason so many genes remain cryptic in their function is because this system is frequently studied in the laboratory, removed from its natural context (Petersen et al. 2015), rotting plant detritus in the wild (Kiontke et al. 2011; Frézal and Félix 2015). In the past decade, the natural microbial environment of *C. elegans* has been characterized (Berg, Stenuit, et al. 2016; Dirksen et al. 2016; Samuel et al. 2016; Zhang et al. 2017), and this information could be used to enable laboratory experiments aimed at discovering novel gene functions (Singh and Luallen 2024). Here, we aim to describe the natural microbial environment of *C. inopinata* to (1) harness the fig microcosm to understand the ecological drivers of organismal occupancy and diversity and (2) provide a comparative context to enable laboratory and field studies aimed at understanding how host/microbe interactions evolve. To this end, we performed 16S microbial metabarcoding on fresh *F. septica* figs with variable nematode occupancy and pollinating foundress wasp number to understand whether and how they drive patterns of microbial diversity.

## Results

### 16S metabarcoding reveals bacteria associated with F. septica figs

To describe the microbial taxa associated with *C. inopinata*, we performed 16S metabarcoding (see methods) on substrates from this species’ natural environment. 38 figs from 12 *F. septica* plants in Taipei, Taiwan were sampled in August 2019 (Figure 1). Two different DNA preparations were generated for each fig: (1) before dissection, the fig was washed in buffer and this surface wash was then used for downstream sequencing (known herein as “fig surface wash” samples); (2) the fig was then cut into four pieces in buffer, and a fraction of the resultant fig suspension was used for downstream sequencing (known herein as “fig suspension” samples). In addition to microbial metabarcoding, additional observations were collected for each dissected fig including nematode occupancy and foundress pollinating wasp number (see Supplemental Table 1 sheet 3 for all fig-specific ecological data). This enabled the testing of hypotheses regarding the impacts of nematodes and wasps on fig microbial diversity. 16S libraries from these samples were prepped and sequenced, resulting in 11,361,994 250 bp paired-end reads (140,272 paired-end reads on average per sample; Supplemental Figure 1; Supplemental Table 1 sheet 2). Reads were then clustered to infer Operational Taxonomic Units (OTUs); 9,421,644 paired-end reads were ultimately used to infer microbial communities (see methods; 116,317 paired-end reads on average per sample; Supplemental Figure 1). An initial analysis revealed the reads were dominated by organellar DNA, presumably originating from host tissue (37% reads mitochondrial or chloroplast in origin on average per sample; per-sample range 0-94%; Supplemental Figure 2). After excluding these reads, 5,215,832 paired-end reads remained for downstream analyses (64,393 paired-end reads on average per sample; Supplemental Figure 1).

**Figure 1.**
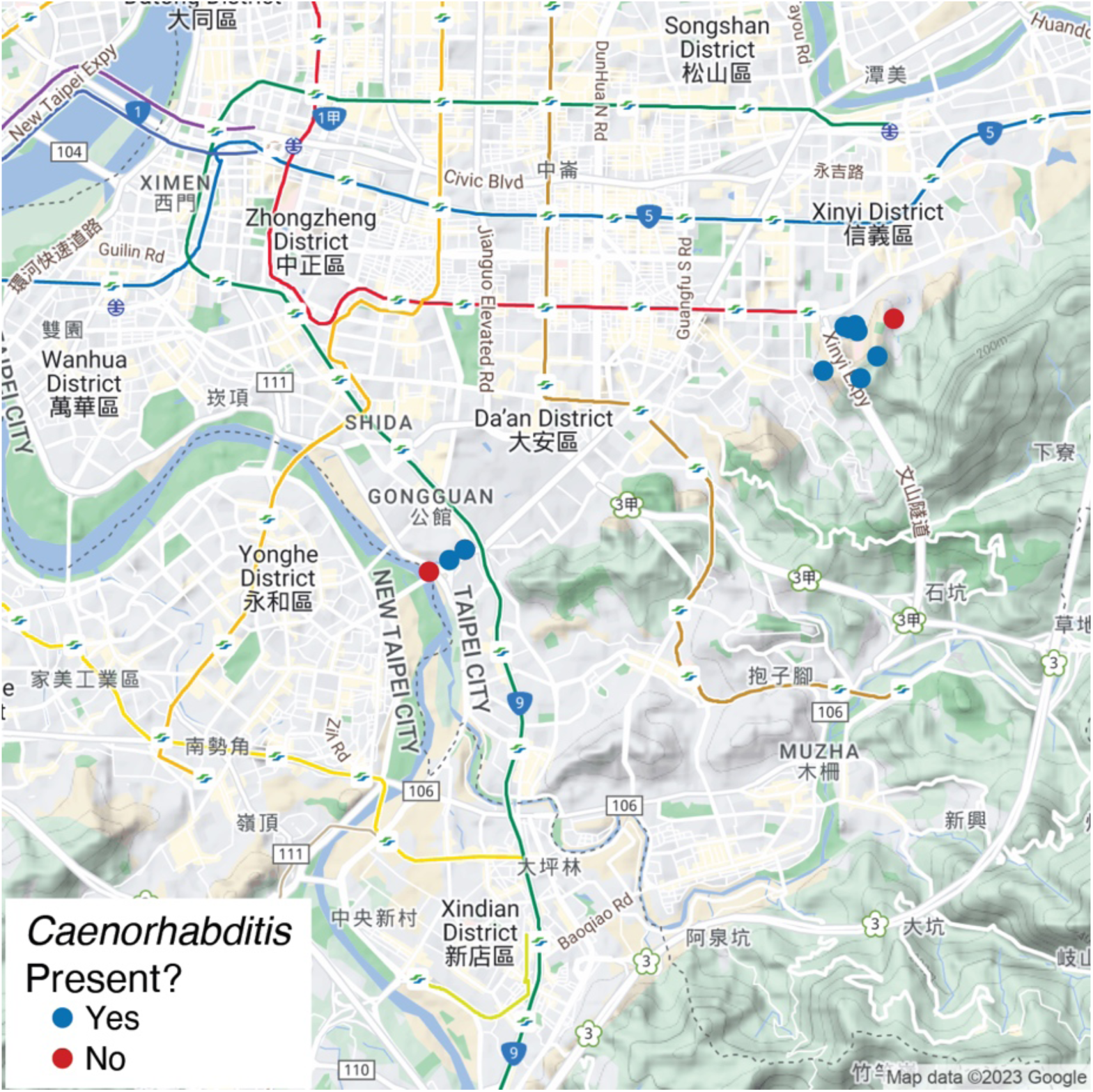
Sampling localities. *F. septica* figs were sampled in southern Taipei, Taiwan. Points represent the locations of individual *F. septica* plants sampled in this study. Points are colored by whether or not any figs observed from a given plant harbored *Caenorhabditis* nematodes. Maps were generated in part with Google Maps and their data sources. © 2023 Google (https://about.google/brand-resource-center/products-and-services/geo-guidelines/).

Highly prevalent taxa are frequently described as constituting a “core” microbiome for a given host (Neu et al. 2021). For an initial description of *Ficus septica* microbial communities, we examined the most prevalent taxa in our samples. Here, no OTUs were observed in 100% of our fig samples (Supplemental Figure 3), but one genus was observed in all fig samples, *Methylobacterium/Methylorubrum* (Green and Ardley 2018). As these OTU- and genus-level taxonomic ranks typically revealed lower prevalence (Supplemental Figures 3-5), we proceeded to describe fig microbial diversity at the family level (Figure 2). Twenty bacterial families were observed in at least 80% of fig samples (which we refer to here as “high prevalence”) when either considering all samples together or within-group prevalence in either fig suspensions or fig surface washes alone (Figure 2). These families are nested within the taxonomic classes Actinobacteria, Bacteroidia, Alphaproteobacteria, and Gammaproteobacteria (Figure 2). Eleven families revealed high prevalence in all samples (including *Acetobacteraceae*, *Beijerinckiaceae*, *Moraxellaceae*, *Spirosomaceae*, and *Kineosporiaceae*; Figure 2). Of the nine remaining families, eight had high prevalence in fig suspensions but <80% prevalence in fig surface washes (*Nocardiaceae*, *Chitinophagaceae*, *Sphingomonadaceae*, *Weeksellaceae*, *Rhodobacteraceae*, *Enterobacteriaceae*, and *Erwiniaceae*; Figure 2). In contrast, the last family (*Geodermatophilaceae*) had high prevalence in fig surface wash samples but only 76% prevalence in fig suspension samples (Figure 2). Ultimately, 3,321 OTUs, 306 genera, and 182 families were identified in our fig samples. Most OTUs exhibited low prevalence (mean prevalence 3%; Supplemental Figure 3). Broadly, fig OTUs were largely associated with the Proteobacteria (37.0% of OTUs), Bacteroidota (19.8%), and Actinobacteriota (9.9%) phyla. This suggests *F. septica* harbors a Proteobacteria-rich microbiota, typical of those associated with plants (Müller et al. 2016).

**Figure 2.**
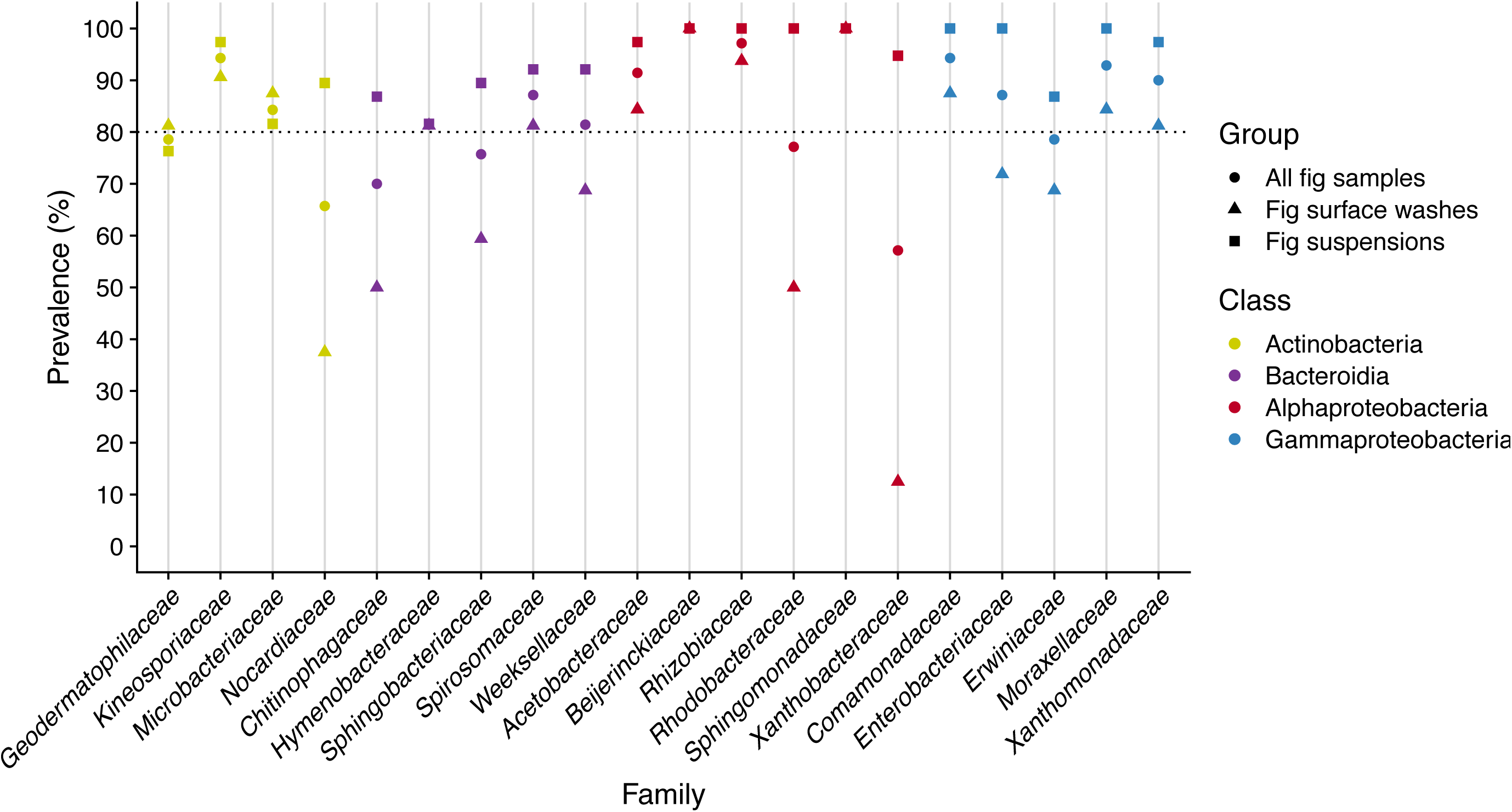
Bacterial family prevalence in fig samples. Families prevalent in 80% of the samples (with respect to a given subset of the samples: all fig samples, fig surface washes, or fig suspensions) were plotted. Note, in the Silva taxonomy, the family *Spirosomataceae* is noted as *Spirosomaceae*. In addition, in this taxonomy, *Comamonadaceae* is noted as a member of the class Gammaproteobacteria despite its status as a member of Betaproteobacteria in other sources (Yilmaz et al. 2014).

*F. septica* figs are where *C. inopinata* nematodes thrive (Kanzaki et al. 2018; Woodruff and Phillips 2018). As *C. inopinata* is the sister species of the *C. elegans* model system (Kanzaki et al. 2018), the definition of highly-prevalent taxa in *F. septica* figs affords a comparison with microbial communities associated with the natural environment of *C. elegans*. Thirteen families were found to be present in 100% of samples associated with *C. elegans* (Samuel et al. 2016; Zhang et al. 2017). When compared with the twenty high prevalence families detected in our study (Figure 2), ten of these families were shared (Supplemental Figure 6; Supplemental Table 1 sheet 4). The other ten families were not fig-specific; rather, they were found in lower prevalence in *C. elegans* samples (Supplemental Table 1 sheet 4). Three families defined as highly-prevalent in *C. elegans* samples (Zhang et al. 2017) did not overlap with our high prevalence fig families (Supplemental Figure 6; Supplemental Table 1 sheet 4). These families were not *C. elegans*-specific; rather, they were found at somewhat lower prevalence among our fig samples (37-67%; Supplemental Figure 6; Supplemental Table 1 sheet 4). Thus, *C. inopinata* and *C. elegans* bacterial communities largely overlap but vary in their prevalence.

### Fig exteriors and fig interiors harbor distinct microbial communities

Interior and exterior fig microbial communities are expected to be distinct because only fig interiors harbor pollinating wasps and their associated nematodes (and interior communities are expected to harbor microbes directly associated with *C. inopinata*). Here, we use fig surface wash samples to capture exterior communities, and we use fig suspension samples to capture interior communities (although such samples will also likely include taxa from fig surfaces as well). The prevalence metrics discussed above suggest differences among the communities of surface washes and fig suspensions (Figure 2). Consistent with this, a Bray-Curtis ordination of samples reveals the separation of fig surface wash and fig suspension samples across major axes of variation (Figure 3, Supplemental Figures 7 & 9). However, this pattern varies depending on the distance metrics used, with unweighted UniFrac distances revealing little separation of fig suspension and fig surface wash samples following ordination (Supplemental Figures 8, 10-11). Nonetheless, microbial communities among fig surface washes and fig suspensions are distinct under a hypothesis testing framework (PERMANOVA *p* = 0.001; *F* = 3.2; pseudo-*r^2^* = 0.045; Supplemental Table 1 sheet 5). Comparisons of OTU abundances reveal 11 OTUs to have significant differences in abundance between fig surface washes and fig suspensions (False discovery rate-adjusted Wilcoxon rank-sum test *p* < 0.05); Supplemental Figure 12; Supplemental Table 1 sheet 6). Consistent with this, 15 genera harbored significant differences among surface washes and suspensions (False discovery rate-adjusted Wilcoxon rank-sum test *p* < 0.05; Supplemental Figure 13; Supplemental Table 1 sheet 7). Correspondingly, 15 families similarly revealed significant differences among groups (False discovery rate-adjusted Wilcoxon rank-sum test *p* < 0.001); Supplemental Figure 14; Supplemental Table 1 sheet 8). Families enriched on fig surface washes included *Beijerinckiaceae*, *Microbacteriaceae*, and *Sphingomonadaceae* (Cohen’s *d* effect size range = −1.42 — −0.78; Supplemental Figure 14; Supplemental Table 1 sheet 8); families enriched in fig suspensions included *Xanthobacteraceae*, *Rhodobacteraceae*, and *Moraxellaceae* (Cohen’s *d* effect size range = 0.92-2.26; Supplemental Figure 14; Supplemental Table 1 sheet 8). However, fig surface washes and fig suspensions revealed no differences in within-sample α-diversity across three measures (Supplemental Figure 15; Wilcoxon rank-sum test *p* = 0.28 (Number of OTU), *p* = 0.26 (Shannon diversity), *p* = 0.10 (Faith’s phylogenetic diversity)). Taken together, microbial communities appear distinct between fig exteriors and fig interiors.

**Figure 3.**
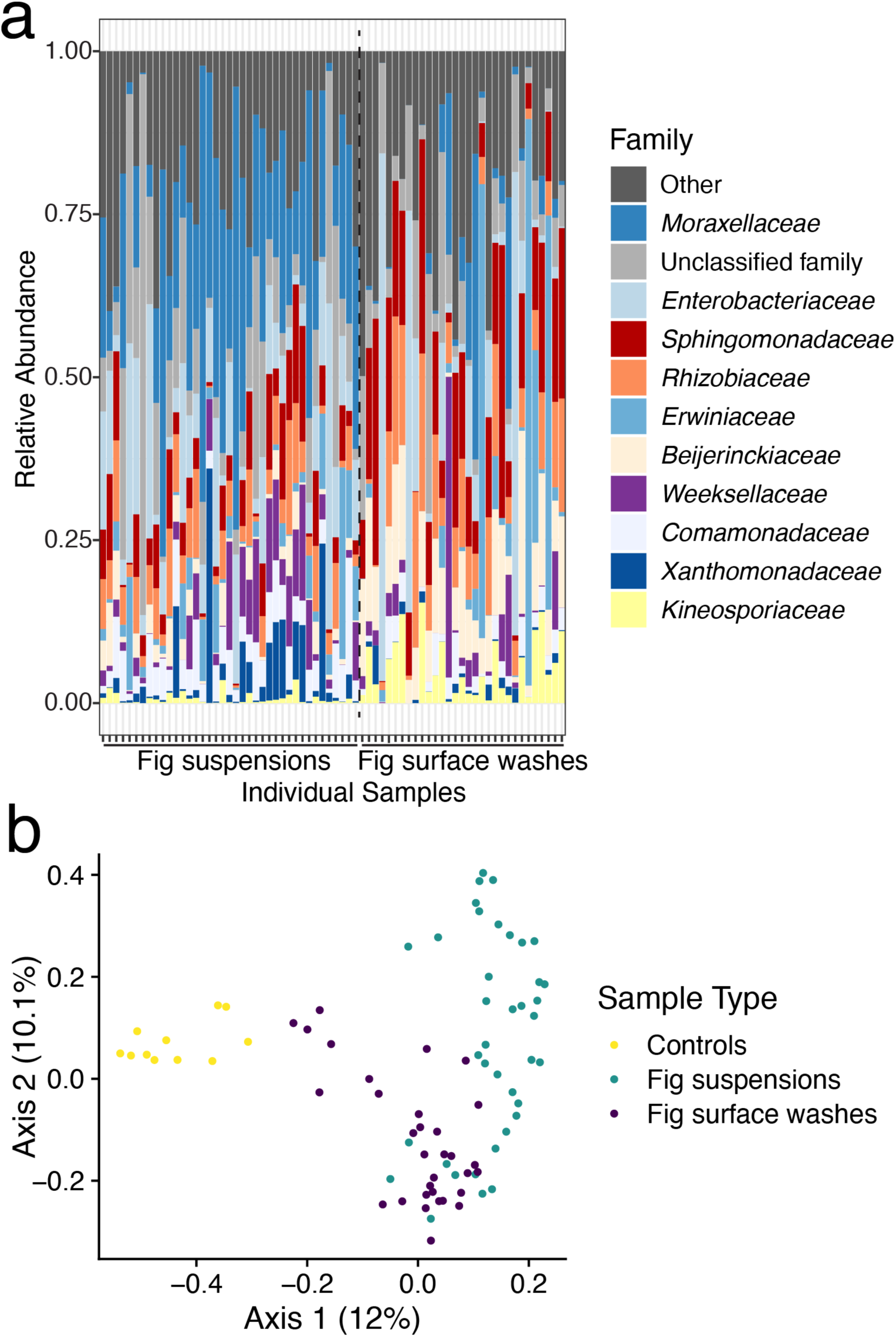
The microbial communities of fig surfaces and fig homogenates can be distinguished. (a) A composition plot reveals the relative abundance of microbial families across samples. Samples are ordered along the *x*-axis by sample type (fig suspensions on the left; fig surface washes on the right. They are separated by the vertical dotted line.). The *y*-axis notes the relative proportion of reads corresponding to a given family in the samples. (b) Principal Coordinate Analysis using Bray-Curtis dissimilarity distances reveals distance gradients among sample types. Each point represents a sample. The major axis separates control samples from fig samples, and the second axis separates fig homogenates from fig surface washes.

### Nematodes have a modest impact on fig microbial communities

To understand the impact of fig nematodes on fig microbial communities, metabarcoding data were connected to measures of nematode occupancy made when figs were initially sampled. Here, we focused on internal fig suspensions as *C. inopinata* nematodes thrive inside figs (Woodruff and Phillips 2018). Nematode occupancy does have a modest impact on microbial communities (Figure 4a; PERMANOVA *p* = 0.04; *F* = 1.3; pseudo-*r^2^* = 0.035; Supplemental Table 1 sheet 5). No OTUs (or other taxa regardless of rank) revealed significant differential abundance among figs with and without nematodes (after correcting for multiple comparisons; all false discovery rate-adjusted Wilcoxon rank-sum test *p* > 0.05; Supplemental Table 1 sheets 9-14). However, 17 OTU did reveal differential abundance in the absence of corrections for multiple comparisons (Supplemental Figure 16; uncorrected Wilcoxon rank-sum test *p* < 0.05; Supplemental Table 1 sheet 9). Consistent with these observations, samples varying in nematode occupancy overlapped in ordination space (Figure 4b; Supplemental Figures 17-19) Additionally, nematode occupancy revealed little impact on within-sample α-diversity (Figure 5; Wilcoxon rank-sum test *p* = 0.38 (Number of OTU), 0.50 (Shannon diversity), 0.46 (Faith’s phylogenetic diversity)). Thus, while nematodes may have some influence on fig microbial composition, they appear to promote little observable impact on microbial α-diversity.

**Figure 4.**
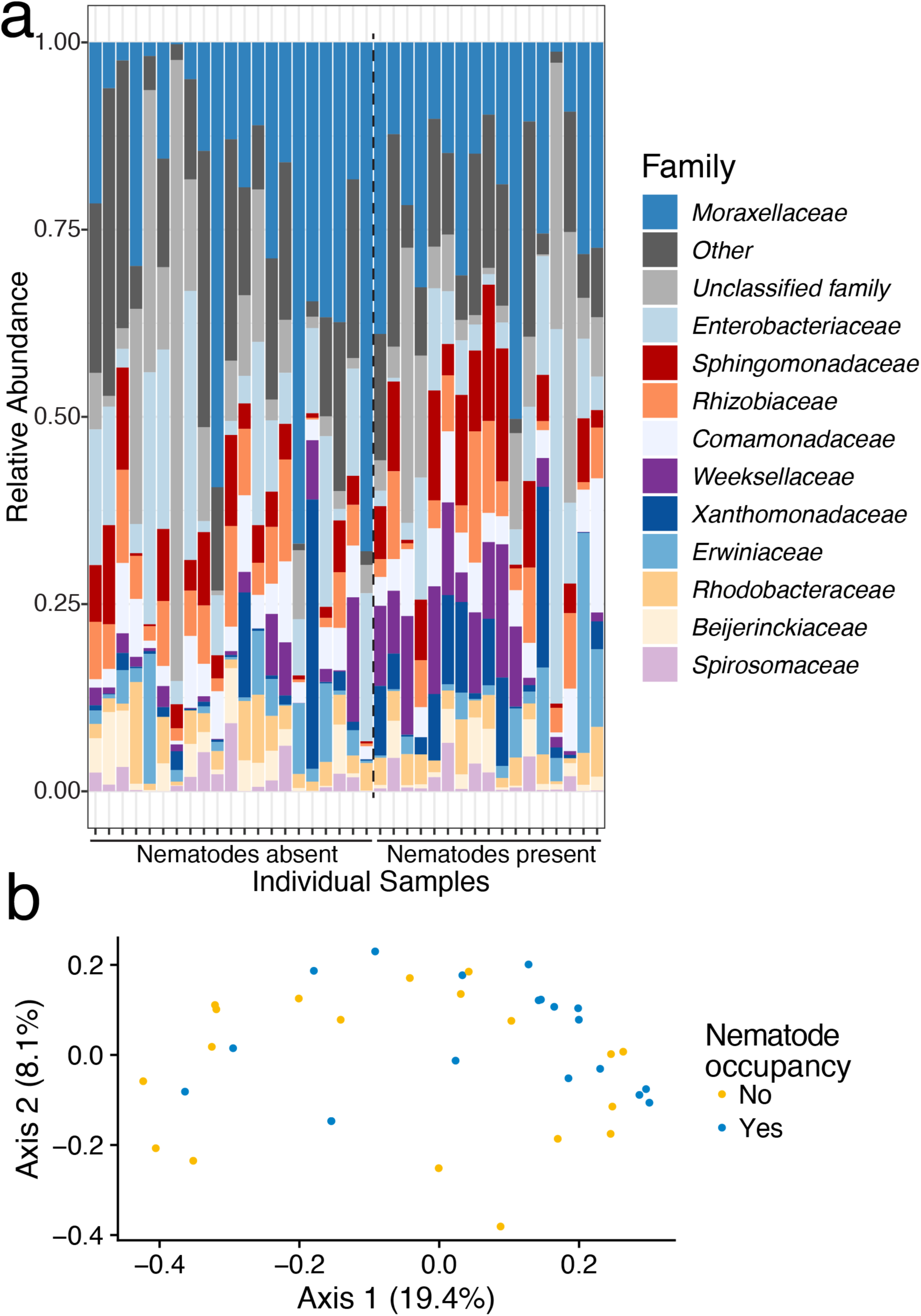
Nematodes have a modest impact on fig microbial communities. Here, only fig suspension samples are considered. (a) A composition plot reveals the relative abundance of microbial families across samples. Samples with nematodes present, left; samples without nematodes, right (separated by the vertical dotted line). The *y*-axis notes the relative proportion of reads corresponding to a given family in the samples. Note, in the Silva taxonomy, the family *Spirosomataceae* is noted as *Spirosomaceae*. (b) Principal Coordinate Analysis using Bray-Curtis dissimilarity distances reveals distance gradients among sample types. Each point represents a sample.

**Figure 5.**
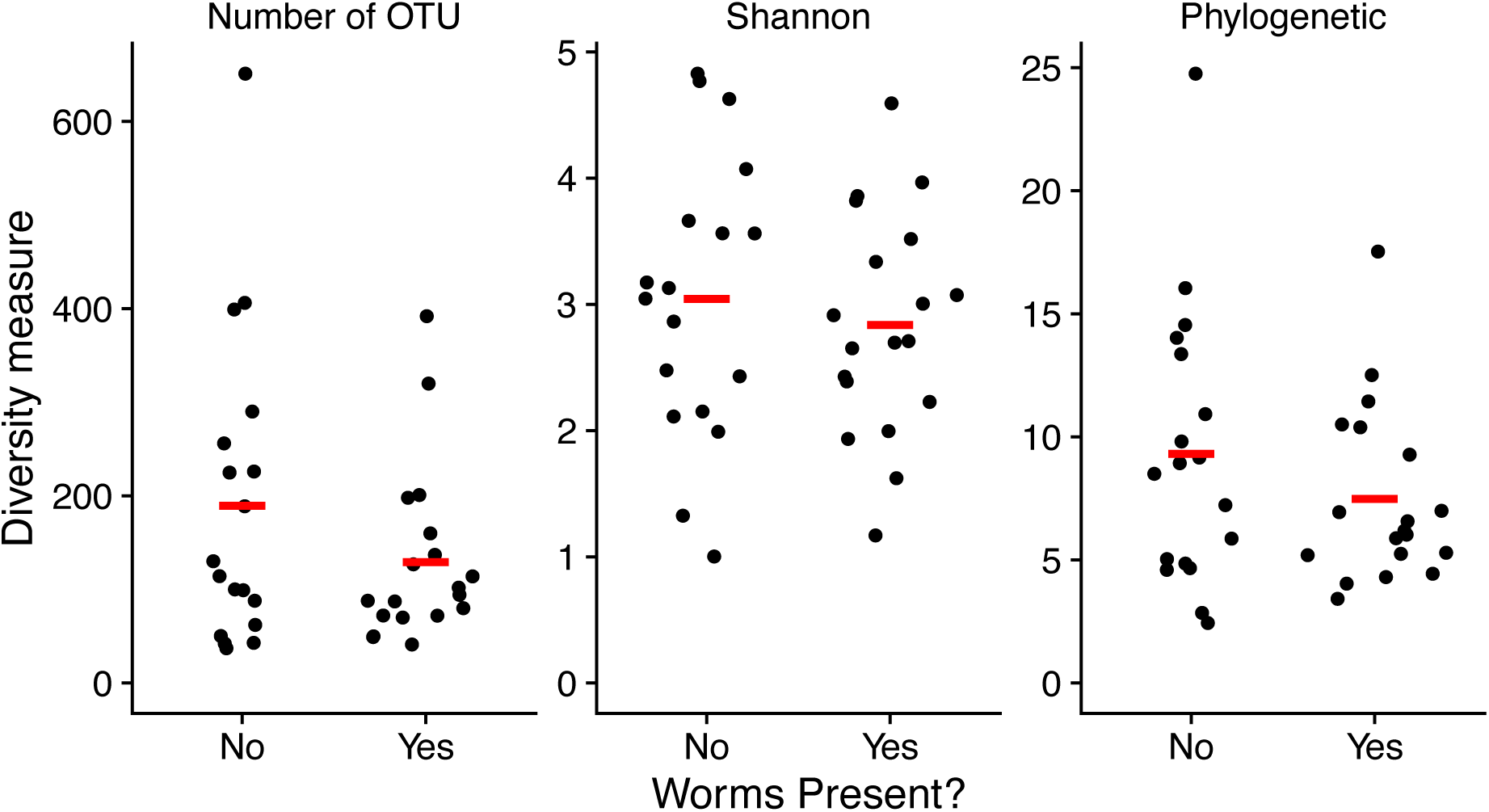
Nematodes do not influence fig microbial α-diversity. Here, only fig suspension samples are considered. Three diversity metrics (number of OTU’s found in each sample; Shannon diversity; and Phylogenetic diversity) are plotted for samples with or without nematodes. Sina plots are strip charts with points taking the contours of a violin plot. Horizontal bars represent means. None of these comparisons are statistically significant (Wilcoxon rank-sum test *p* > 0.05).

### Fig microbial diversity is negatively correlated with foundress wasp number

Figs can be pollinated by more than one wasp, and foundress number can have profound influences on the fig microenvironment. For instance, as foundress number increases, wasp progeny sex ratios become more biased (Herre 1985; Herre 1987; Molbo et al. 2004), presumably due to increased local mate competition (Hamilton 1967). In addition, as foundress number increases, the probability of fig nematode occupancy increases (Woodruff and Phillips 2018), likely because nematodes travel on foundress pollinators (Woodruff and Phillips 2018; Shi et al. 2019). Because of these effects of foundress number on fig biology, it was natural to suspect this could also impact fig microbial diversity. Indeed, foundress number is negatively correlated with within-sample phylogenetic α-diversity (Figure 6A; Faith’s phylogenetic diversity; ordinary least squares regression *p*= 0.015, *F*= 6.6, *r^2^*= 0.16)). However, for two other metrics of α-diversity, no significant relationships between within-sample α-diversity and foundress number were detected (Figure 6A; ordinary least squares regression for Number of OTU: *p*= 0.23, *F*= 5.8, *r^2^*= 0.14; Shannon diversity: *p*= 0.11, *F*= 2.7, *r^2^*= 0.044). Additionally, samples with high pollinator burdens tend to cluster in ordination space (Figure 6B; Supplemental figures 20-22). Thus, while nematode occupancy has a seemingly subtle impact on fig microbial diversity, wasp pollinators appear to have a larger influence on these communities.

**Figure 6.**
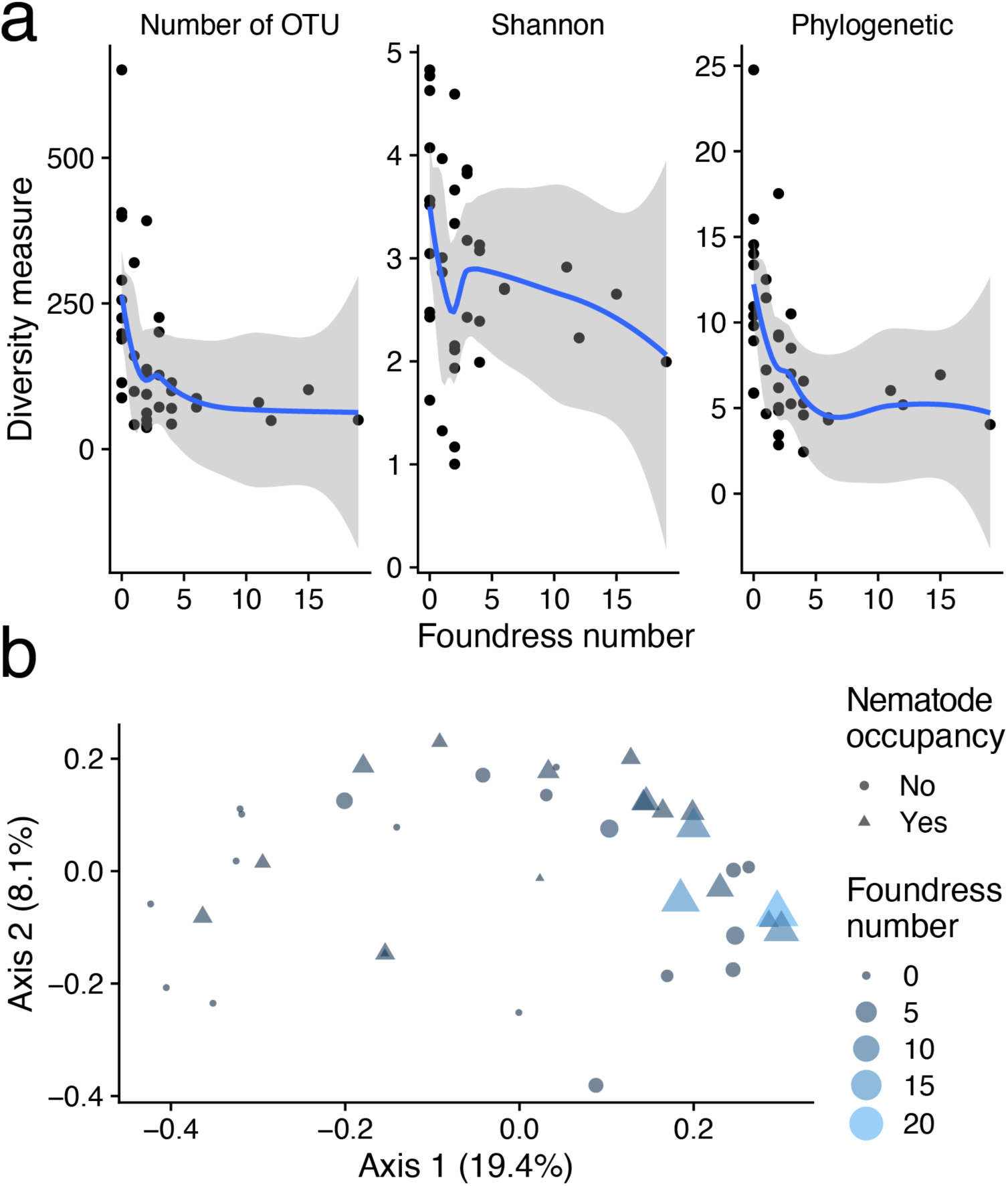
Fig microbial phylogenetic diversity is negatively correlated with the number of foundress wasps. Here, only fig homogenate samples are considered. (a) Three diversity metrics (number of OTU’s found in each sample; Shannon diversity; and Phylogenetic diversity) are plotted against the pollinating wasp foundress number associated with that fig sample (linear model *p* < 0.05 for Phylogenetic diversity; *p* > 0.05 for Number of OTU and Shannon diversity; other linear model statistics are in the main text). The solid lines were generated with LOESS regression; the shaded area represents 95% confidence intervals. (b) Principal Coordinate Analysis using Bray-Curtis dissimilarity distances reveals distance gradients among sample types. Each point represents a sample. This is the same plot as in Fig. 4b, but with pollinator foundress wasp number added as an additional dimension (point size and shade).

## Discussion

### The microbial community of Ficus septica figs

Here, we used high-throughput metabarcoding to understand the microbial communities associated with *Ficus septica* figs. We identified over 3,000 OTUs across over 300 genera and nearly 200 families. Among these taxonomic groups, *Methylobacterium/Methylorubrum* species were found in all of our fig samples. Notable for being able to grow on single-carbon substrates such as methanol or methylamine, these species have been frequently isolated from plants (Holland and Polacco 1994; Green and Ardley 2018). Moreover, *Methylobacterium/Methylorubrum* species have been shown to be beneficial to plant hosts— *Methylobacterium mesophilicum* can rescue urease mutants in soybean (Holland and Polacco 1994), and certain strains of *Methylobacterium extorquens* can increase the germination rate of sugarcane (Madhaiyan et al. 2005). *Sphingomonas* was another highly-prevalent *Alphaproteobacteria* species in our fig samples. *Sphingomonas* species are likewise commonly found in plant substrates (Asaf et al. 2020) and also promotes growth in tomatoes (Khan et al. 2014) and *Arabidopsis* (Luo et al. 2019). *Kineococcus* species were also abundant in our samples (Supplemental Figure 5); these *Actinobacteria* have also been isolated from diverse substrates including soil (Lewin et al. 2016), desert sand (Lewin et al. 2016), angiosperm roots (Duangmal et al. 2008), and human-built environments (Mhatre et al. 2020). *Acinetobacter* species were present in all fig suspension samples (Supplemental Figure 5). Notable because of their capability to promote opportunistic infection in humans, these bacteria have also been isolated from diverse environments, including soil, water, and arthropods (Doughari et al. 2011; Jung and Park 2015). Other relatively dominant genera in our samples (such as *Spirosoma* (Finster et al. 2009) and *Chryseobacterium* (Jung et al. 2023)) have also been isolated from a diverse array of similar environments like soil, roots, and other plant-associated substrates. Taken together, the microbial community of *Ficus septica* harbors taxa that have been previously shown to be plant-associated (some with demonstrated plant-beneficial effects) or have been otherwise isolated from diverse habitats.

### The microbes of Ficus species and their associated wasps

Other studies have characterized fig microbial diversity— how do our results compare? While there is some overlap with other studies, the microbial communities of *Ficus septica* appear largely divergent from those reported in *Ficus hirta* and *Certaosolen* and *Eupristina* fig wasps. Liu et al. examined male and female *Ficus hirta* syconia; their samples were dominated by *Wolbachia*, *Ralstonia*, and *Burkholderia* species (Liu et al. 2021). Here, only two samples harbored *Wolbachia*, and no samples harbored *Ralstonia* or *Burkholderia*. Moreover, in our study, 11 (out of 12) male figs revealed no *Wolbachia* OTUs. It is unclear why so few *Wolbachia* reads were found in our samples. Future work directly characterizing pollinating *Ceratosolen* and parasitic *Philotrypesis* wasps associated with *F. septica* figs (Woodruff and Phillips 2018)) themselves would be more likely to discover wasp-associated endosymbionts. Two other studies examined the bacterial communities associated with fig wasps. A study examining *Ceratosolen* fig wasps (including those associated with *Ficus hispida*, known to harbor *Caenorhabditis* nematodes (Jauharlina et al. 2022)) reported samples dominated by *Proteobacteria* (including both Gammaproteobacteria and Alphaproteobacteria), *Firmicutes*, and *Bacteroides* (Li et al. 2022). Another study examined *Eupristina* fig wasps associated with *Ficus altissima* and *Ficus microcarpa*; here, the dominant genera included *Propionibacterium*, *Streptococcus*, *Acinetobacter*, *Staphylococcus*, and *Klebsiella* (Dong et al. 2022). Two of these (*Acinetobacter* and *Klebsiella*) were also common in our samples. Additionaly, these other studies do not report many of the most common taxa in our samples, including *Methylobacterium*, *Allorhizobium*, and *Sphingomonas*. Additionally, although other studies have looked at the fungal communities of figs (Martinson et al. 2012; Dong et al. 2022), our data cannot speak to the extent and constituents of fungal diversity in these samples. Regardless, it is then possible there is great diversity in the microbial communities across *Ficus* fig microcosms.

### The microbes of *C. inopinata* and *C. elegans*

A major goal of this work was to discover the microbes associated with the bacterivore *C. inopinata* to enable comparisons with the *C. elegans* model system. However, the dynamic pace of microbial taxonomy in tandem with variation in taxonomic schemes used by differing metabarcoding studies hampers comparisons. That is, apparent differences in microbial communities may simply represent differences in taxonomic systems (or changes in taxonomic conventions over time). Despite these caveats, there was overlap among the taxa found in *F. septica* figs and those commonly associated with *C. elegans* (Supplemental Figure 6) (Berg, Stenuit, et al. 2016; Dirksen et al. 2016; Samuel et al. 2016; Zhang et al. 2017). Like *C. elegans*, substrates associated with *C. inopinata* were rich in *Proteobacteria* and were associated with OTUs in the *Actinobacteria* and *Bacteroides* phyla (Figures 2-4). More specifically, out of 14 families reported across all natural *C. elegans* microbiomes (Zhang et al. 2017), ten overlapped with families found to be highly prevalent (>80%) among some subset of our fig samples (*Microbacteriaceae*, *Comamonadaceae*, *Enterobacteriaceae*, *Moraxellaceae*, *Xanthomonadaceae*, *Weeksellaceae*, *Acetobacteraceae*, *Sphingomonadaceae*, *Sphingobacteriaceae*, and *Rhodobacteraceae*; Supplemental Figure 6). However, some genera frequently found in our samples did not overlap with the top 260 common OTUs found in *C. elegans* (such as *Kineococcus*, *Rhizobium*, *Quadrisphaera*, and *Spirosoma*; Supplemental Figure 5; see Supplemental Table 2 of (Zhang et al. 2017)). And, some families with high prevalence in *C. elegans* samples were not as common among our samples (*Pseudomonadaceae*, *Oxalobacteraceae*, and *Flavobacteriaceae*). Taken together, these seemingly radically divergent ecological niches (rotting plant detritus and fresh figs) reveal surprising overlap in their microbial communities while harboring clear differences in their most prevalent constituents. Previous studies have noted divergent microbial communities associated with different *Caenorhabditis* species (Berg, Zhou, et al. 2016; Dirksen et al. 2016), suggestive of species-specific ecological niches. The divergent *C. inopinata*-associated microbial community is consistent with the notion that microbial environments are important drivers of species divergence. However, vastly more ecological work is required to determine whether *Caenorhabditis* species occupancy and divergence is driven by the composition of microbial environments. Additionally, other studies have examined the microbial communities of nematodes themselves (apart from their associated substrates (Dirksen et al. 2016)). Here, we only examined substrates (either fig suspensions or surface washes), so we are unable to disentangle nematode microbes from fig microbes *per se*. Future work examining nematodes isolated from their environmental sources will be able to identify such taxa.

### The drivers of fig microbial diversity

Here, we showed that the microbial communities of fig interiors (suspensions) were distinct from those of fig exteriors (surface washes; Figure 3). As fig nematodes thrive in the lumen of the fig (Woodruff and Phillips 2018), they are likely not to be associated with taxa that are restricted to the fig surface. Thus, taxa such as these might be less likely to interact with species like *C. inopinata*. However, we detected no unique genera to the fig surface, although some were enriched in abundance on the fig surface relative to fig suspensions (Supplemental Figure 13). In addition, no differences in microbial diversity among fig suspensions and fig surface washes were detected (Supplemental Figure 15). If anything, there is a trend of *increased* microbial diversity in fig suspensions compared to fig surface washes (Supplemental Figure 15). This was somewhat unexpected, as the fig lumen has strong barriers to entry (Castro-Cárdenas et al. 2022), and we reasoned that as the fig interiors were less exposed, they might have less complex microbiomes. As this was not the case, it is possible that simply the increased biotic material in these samples (compared to surface washes) contributed to this potentially increased microbial diversity. Along these lines, fig suspensions contained nematode, wasp, and fig tissue that was not present in the fig surface wash samples. This is likely the most parsimonious explanation of the diversity observed in fig interiors.

Additionally, we observed a trend of declining microbial diversity with foundress wasp number (Figure 6). Foundress wasps are known to change the fig environment—they introduce pollen, promoting seed development, and they lay eggs in fig ovules. And, they carry nematodes and microbes that carry the potential to influence the fig environment. Indeed, the probability of nematode occupancy increases as foundress number increases in these figs (Woodruff and Phillips 2018). So, it is perhaps unsurprising that the microbial community likewise changes with foundress number. It is unclear why microbial diversity should *decrease* with foundress number. One possibility is that bacterivorous nematode load increases with foundress number, which leads to a reduction in microbial diversity (due to microbe consumption by nematodes). However, we saw no influence of nematode occupancy on microbial diversity as such (Figure 5). Additionally, foundress wasps may carry bacterial taxa that dominate the fig upon colonization, leading to decreased diversity. Consistent with this, evenness indices likewise exhibit a negative relationship with foundress number (at least via one metric (Smith and Wilson’s Evar index); Supplemental Figure 25). Alternatively, increased pollination, oviposition, or both may induce changes in the fig environment that impact microbial diversity. Regardless, future studies with a larger range of phenotypic conditions, in addition to manipulative field experiments, will be needed to disentangle these possibilities.

We also detected only a modest impact of nematode occupancy on microbial communities (Figure 4), with no detectable impact on microbial diversity as such (Figure 5). This is surprising because *C. elegans* occupancy has been shown to have an influence on substrate communities (Samuel et al. 2016), with nematode abundance promoting a reduction of diversity (and with reproductive and dispersal stage-biased populations harboring differing microbial communities (Samuel et al. 2016)). It is unclear why no influence was observed— one possibility is that *C. inopinata*, despite its bacterivorous status, simply does not have a large effect on fig microbial communities. This may particularly be true if *C. inopinata* does not eat much bacteria, but rather more fungi, protists, or other particles in the fig. Indeed, the natural food source of *C. inopinata* has not yet been determined. However, it is also possible that we do not yet have the power to detect the impact of *C. inopinata* on fig microbial communities, and that with more figs, more reads, or a more dynamic range of fig conditions (i.e., figs with a greater range in nematode loads or nematode stages) would reveal greater variation in fig microbial communities. Despite this, we did potentially detect some OTU that may be associated with nematode occupancy (Supplemental Figure 16). Ongoing metagenomic work on individual nematodes from the field, as well as ongoing experimental work with fig-derived microbes, will be invaluable towards informing these possibilities.

### Implications for the biology of *C. inopinata*

*C. inopinata* is an ecologically and morphologically exceptional *Caenorhabditis* species (Kanzaki et al. 2018; Woodruff et al. 2018; Woodruff and Phillips 2018). In a genus largely characterized by profound phenotypic stasis across large genetic distances (e.g., *C. briggsae* and *C. elegans* are nearly morphologically indistinguishable despite harboring genetic divergence comparable to that of human and mouse (Kiontke et al. 2004)), the current sister species of *C. elegans* is an outlier. *C. inopinata* is large in size, grows slowly, and thrives in a novel environment (Kanzaki et al. 2018; Woodruff et al. 2018; Woodruff and Phillips 2018; Woodruff et al. 2019). Here, we set out to better understand this microenvironment to better inform how its ecological context may influence its phenotypic divergence. While nematode occupancy did not modulate fig microbial diversity as such (Figure 5), there was a significant, but modest, impact on fig microbial composition. Indeed, OTUs were detected that were both enriched (e.g., OTUs from the genera *Ochobactrum*, *Kosakonia*, and *Stentotrophomonas*; Supplemental Figure 16) and depleted (e.g., OTUs from the genera *Acinetobacter*, *Chitinophagaceae*, and *Auriemonas*; Supplemental Figure 16). Thus, these taxa may interact with *C. inopinata* in some way—either as food, pathogens, or otherwise beneficial or detrimental species. At least five of these taxa have been previously reported in *C. elegans* samples (Samuel et al. 2016; Zhang et al. 2017), and three of them have been included in the CeMbio resource, a simplified, standardized natural *C. elegans* microbiota for experimental studies (Dirksen et al. 2020). This supports the notion these taxa represent the natural food of (or may otherwise interact with) *C. inopinata*. Alternatively, these patterns may emerge via indirect consequences of nematode occupancy via their potential impacts on other community members. Or, these presumptive nematode-responsive taxa may represent false positives. Additionally, we also detected a decline of microbial diversity with foundress wasp number (Figure 6). As the probability of worm occupancy increases with foundress number (as nematodes travel on wasps), it is reasonable to suspect that within-fig nematode population sizes increase as foundress number increases. Thus, nematodes may be more likely to be exposed to microbial communities associated with crowded figs. Nematodes may experience more frequent (or stronger) selection in these environments, and microbial taxa enriched in high foundress number figs may represent better proxies for a core microbial community of *C. inopinata*. For instance, *Enterobacterales* bacteria were highly abundant in a fig with 19 foundresses; this may be a family that interacts more frequently with *C. inopinata*. Indeed, microbes frequently associated with *C. elegans* in nature can modulate fitness-related traits in laboratory conditions (Dirksen et al. 2016; Samuel et al. 2016). Continued sampling of high foundress number figs; the sequencing of individual *C. inopinata* nematodes isolated from natural environments; together with targeted microbial culture strategies will build out the resources required to understand the novel environment of *C. inopinata* and how it influences its divergent phenotypes.

### Caveats

Limitations to this work exist. As mentioned above, although the goal of this study is to characterize the microbial communities associated with *C. inopinata*, it cannot delineate the *C. inopinata* microbiome as such. As we used whole fig suspensions, we cannot distinguish between microbes that directly interact with *C. inopinata* from those that do not. Indeed, many figs were also occupied by wasps and fungi, and some were occupied by other animals (such as mites, non-wasp insect larvae, and ants). And, of course, our samples were dominated by fig tissue. Thus, these results are best interpreted as a description of the *F. septica* fig microcommunity, and not one of the microbiomes of a specific fig microcommunity member. To understand nematode microbiomes, we will need to sequence individual nematodes (and, surface-sterilized nematodes have been used to potentially interrogate the gut microbiome (Dirksen et al. 2016; Kanzaki et al. 2018), which is of broad interest). Connected to this is the large number of organellar reads observed in our data (Supplemental Figure 2). Presumably, these are derived from the host and other eukaryotic community members. This suggests that these results may need to be approached with caution, as rarer microbial taxa were perhaps not detected as most reads were covered by uninformative non-bacterial sources. However, a rarefaction analysis reveals that for most samples, coverage was high enough to capture the microbial diversity present (Supplemental Figures 23-24). Yet, future studies using alternative library preparation methods to limit organellar DNA amplification (Lundberg et al. 2013) may prove useful in saving costs due to the generation of such uninformative sequences (and to mitigate any biases these sequences may introduce). Additionally, our experience in analyzing these results leads us to suspect taxonomy has an outsized influence—results and interpretation can vary depending on the taxonomic system implemented. There may be similarities between our work and previous studies that we may have overlooked due to differences in taxonomic system; likewise, there may have been important differences among studies that we may have overlooked for the same reason. Finally, metabarcoding provides little information regarding the biochemical functional roles of microbial community members. Future work assembling whole bacterial genomes associated with these figs will be required to delineate functional roles among community members to understand if they likewise covary with nematode occupancy.

### Concluding thought

Here we described the microbes of the natural environment of the sister species of a key model genetic system. This sets the stage for future work aimed at generating hypotheses regarding the functions of uncharacterized genes and the evolution of ecological divergence. The promise of the *C. inopinata* system lies in the potential to test molecular genetic hypotheses regarding the causes of ecological divergence. To even begin to do this, we must properly describe its divergent environment. That is what we have done here.

## Materials and methods

### Sampling

*Ficus septica* figs were sampled in Taipei, Taiwan on August 27, 2019. Figs were then stored at −4°C. 1-2 days after picking (in the John Wang lab at Academia Sinica), each individual fig was placed in a sterile petri dish and washed with 500 μl of sterile M9 buffer. This surface wash was then transferred to a sterile 1.5 mL Eppendorf tube and stored at −80°C. Then, that same fig was sliced into four pieces in 4 ml of sterile M9 buffer. The fig and suspension were then observed to assess: the approximate fig developmental stage (using the scheme of (Galil and Eisikowitch 1968)); the number of foundress wasps; the presence of *Caenorhabditis* nematodes; the presence of nematodes in general; the presence of other nematode morphotypes; the presence of reproductive stage or dauer-like *Caenorhabditis* nematodes; and the presence of non-pollinating wasps. Then, 500 μl of fig homogenate was then transferred to a sterile 1.5 mL Eppendorf tube and stored at −80°C.

### DNA extraction, library preparation, and sequencing

For DNA extraction, samples (250 μl) were added to a 2 ml screw-cap tube with 750 μl of Qiagen PowerBead Solution, 60 μl of Qiagen SL Solution, and approximately thirty 1.4 mm ceramic beads. The samples were homogenized at 1600 rpm using an MP Biomedicals FastPrep-96 for 2.5 minutes. Samples were then digested overnight (~16 hours) with 10 μl of Proteinase K (20 mg/mL) at 56°C. Samples were extracted using the MagAttract PowerSoil DNA Kit (Qiagen) following the manufacturer’s protocol on a KingFisher Flex (Thermo Scientific) and eluted into 100 μl. The library was prepared using The Earth Microbiome Project 16S primer set (Caporaso et al. 2011) (see Supplemental Table 1 sheet 1 for sample barcode information). PCR was performed with Platinum II Hot-Start 2X PCR Master Mix (Invitrogen). A SimpliAmp thermal cycler (Applied Biosystems) was used with the following cycling conditions: initial denaturation at 94°C for 2 min; 35 cycles of denaturation at 94°C for 15 s, annealing at 54°C for 15 s, extension at 68°C for 7 s; and a final extension step at 68°C for 3 min. PCR amplification was confirmed through visualization on a 2% agarose gel. Samples were then purified and normalized to approximately 1.25 – 1.50 ng/μL using the Just-a-Plate PCR Normalization kit (Charm Biotech). A volume of 10 μL from each sample was pooled before being purified and concentrated using a volume of 1.2X of homemade SPRI beads. The final library was checked for concentration using a Qubit 1X dsDNA High Sensitivity Kit (Thermo Fisher Scientific). Purity and fragment size were confirmed using a Bioanalyzer High Sensitivity DNA Kit (Agilent Technologies). Multiple negative and positive controls were prepared throughout the extraction and library preparation process: three M9 buffer negative controls; two DNA extraction negative controls; one DNA extraction positive control; two PCR negative controls; and two PCR positive controls (ZymoBIOMICS Microbial Community DNA Standard). Paired-end 150-bp reads were then generated with the Illumina MiSeq platform.

### Read processing, OTU assignment, microbial abundance, and statistical inference

Raw sequence data were vetted for quality with FastQC (Andrews 2010). OTUs and their counts were resolved with DADA2 (Callahan et al. 2016) implemented in the Qiime2 package (version 2019.4) (Bolyen et al. 2019). Reads were imported into Qiime2 with *qiime tools import* (options --type ‘SampleData[PairedEndSequencesWithQuality]’ --input-format CasavaOneEightSingleLanePerSampleDirFmt). DADA2 was run within Qiime2 with *qiime dada2 denoise-paired* (options --p-trunc-len-f 180 --p-trunc-len-r 115). Pre-formatted full-length sequences and taxonomic assignments from the SILVA database were retrieved from the Qiime2 website (version 2022.2; https://data.qiime2.org/2022.2/common/silva-138-99-seqs-515-806.qza & https://data.qiime2.org/2022.2/common/silva-138-99-tax-515-806.qza) (Quast et al. 2013; Yilmaz et al. 2014; Ii et al. 2021). Taxonomic identities were assigned to OTU’s with *qiime feature-classifier classify-consensus-vsearch* (with default parameters) (Rognes et al. 2016). Additionally, a phylogenetic tree was generated with representative OTU sequences with *qiime phylogeny align-to-tree-mafft-fasttree* (using default parameters) (Katoh et al. 2002; Price et al. 2009). The feature table, taxonomy, and phylogeny were then converted from the QZA file format to CSV, TSV, and NWK formats (respectively) with *qiime tools export* (using default parameters) for downstream analysis.

Statistical analyses were performed in R (R Core Team) with the aid of the *phyloseq* package (McMurdie and Holmes 2013; Sudarshan et al. 2018 Sep 27). For the characterization and analysis of fig surface wash and fig suspension samples, OTUs associated with mitochondria and chloroplasts were excluded. In addition, OTUs defined in control samples were also excluded from the analysis of these fig samples alone (these OTUs were included in analyses that included controls such as Fig. 2B). For data visualization and ordination, transformed OTU counts were used (a ln(1+*x*) transformation was implemented with the *phyloseq* function *transform_sample_counts*). Principal Coordinate Analyses were performed with Bray-Curtis distances using the *phyloseq* function *ordinate* (with options *method* = “MDS”, *distance* = “bray”). Ordinations with Unweighted Unifrac (*phyloseq* function *ordinate* (with options *method* = “PCoA”, *distance* = “wunifrac”)) and Weighted Unifrac (*phyloseq* function *ordinate* (with options *method* = “PCoA”, *distance* = “unifrac”)) were also performed.

For measures of within-sample alpha diversity, two samples with low non-organellar read counts were removed (samples GW20, GW34), and reads were rarified to 12,252 reads per sample (with the *phyloseq* function *rarefy_even_depth* (with options *rngseed* = 123, *replace* = FALSE). Number of unique OTU’s per sample were inferred with the *phyloseq* function *estimate_richness* (option *measures* = “Observed”). Shannon diversity was estimated with the *phyloseq* function *estimate_richness* (option *measures* = “Shannon”). Faith’s phylogenetic diversity was calculated using the *picante* function *pd* (with default parameters), which sums the branch lengths associated with the subset of the phylogenetic tree (inferred with Qiime2 as described above) associated with a given sample. Mann-Whitney U hypothesis tests were performed with the *wilcox.test* function in base R. All *p*-values were corrected for multiple testing using the *p.adjust* function in base R (options method = “BH”). For statistical hypothesis tests, count data were centered log-ratio transformed using the *transform* function in the *microbiome* package (option “clr”). For PERMANOVA, distance matrices were constructed with the *phyloseq* function *distance* (with options method = “euclidean”). Then, PERMANOVA was performed with the *vegan* function *adonis2* (with the formula (*y* ~ surface_or_interior)) to test differences among fig surface wash and fig homogenate communities. With surface wash samples excluded, PERMANOVA was again performed (with the formula (*y* ~ worms_present)) to test the role of worm occupancy on microbial diversity. All data and code associated with this study have been deposited in Github (https://github.com/gcwoodruff/F_septica_16S_microbial_ecology_2024).

The following R packages were used in this study: *ape* (Paradis and Schliep 2019); *cowplot* (Wilke 2020); *data.table* (Barrett et al. 2024); *dplyr* (Wickham et al. 2023); *DT* (Xie et al. 2024); *effsize* (Torchiano 2016); *ggforce* (Pedersen 2022a); *ggmap* (Kahle and Wickham 2013); *ggplot2* (Wickham 2016); *ggpubr* (Kassambara 2023); *lemon* (Edwards 2020); *microbiome* (Lahti and Shetty 2012 2019); *microbiomeutilities* (Shetty and Lahti 2024); *patchwork* (Pedersen 2022b); *phyloseq* (McMurdie and Holmes 2013); *picante* (Kembel et al. 2010); *RColorBrewer* (Neuwirth 2022); *reshape2* (Wickham 2007); *vegan* (Oksanen et al. 2024); and *UpSetR* (Conway et al. 2017).

## Acknowledgements

We would like to thank Patrick Phillips for supporting this work. We thank Kirsten Nakayama, Nicole Yoneishi, and the University of Hawaiʻi at Mānoa Microbial Genomics and Analytical Laboratory Core for preparing libraries. We thank Graham Wiley and the Oklahoma Medical Research Foundation Clinical Genomics Center for sequencing these libraries. We would like to thank Anna Coleman-Hubert and Erik Johnson for their aid in shipping samples, as well as Jung-Chen Hsu for lab support. This work was supported by funding from the National Institutes of Health Grant to G.C.W. (Grant No. 5F32GM115209-03) and to Patrick Phillips (Grant Nos. R01GM102511, R01AG049396, and R35GM131838). This work was also supported by funding from the National Science Foundation to G.C.W. (Award #2238788). This paper reports data obtained at the University of Hawaiʻi at Mānoa Microbial Genomics and Analytical Laboratory Core, which is supported by an Institutional Development Award (IDeA) from the National Institute of General Medical Sciences of the National Institute of Health under grant number P20GM125508. Some of the computing for this project was performed at the OU Supercomputing Center for Education & Research (OSCER) at the University of Oklahoma. We thank Hayley Lanier for providing comments on an earlier version of this manuscript.

## Data availability

Data and code associated with this study have been deposited in Github (https://github.com/gcwoodruff/F_septica_16S_microbial_ecology_2024). FASTQ files have been submitted to the NCBI Sequence Read Archive (http://www.ncbi.nlm.nih.gov/sra) under the BioProject ID PRJNA1170329.

## Additional files

Supplemental figures and tables can be found at *Journal*.

## Notes

### Competing Interest Statement

The authors have declared no competing interest.

