## Supplemental Figures for "The Bacteria of a Fig Microcommunity"

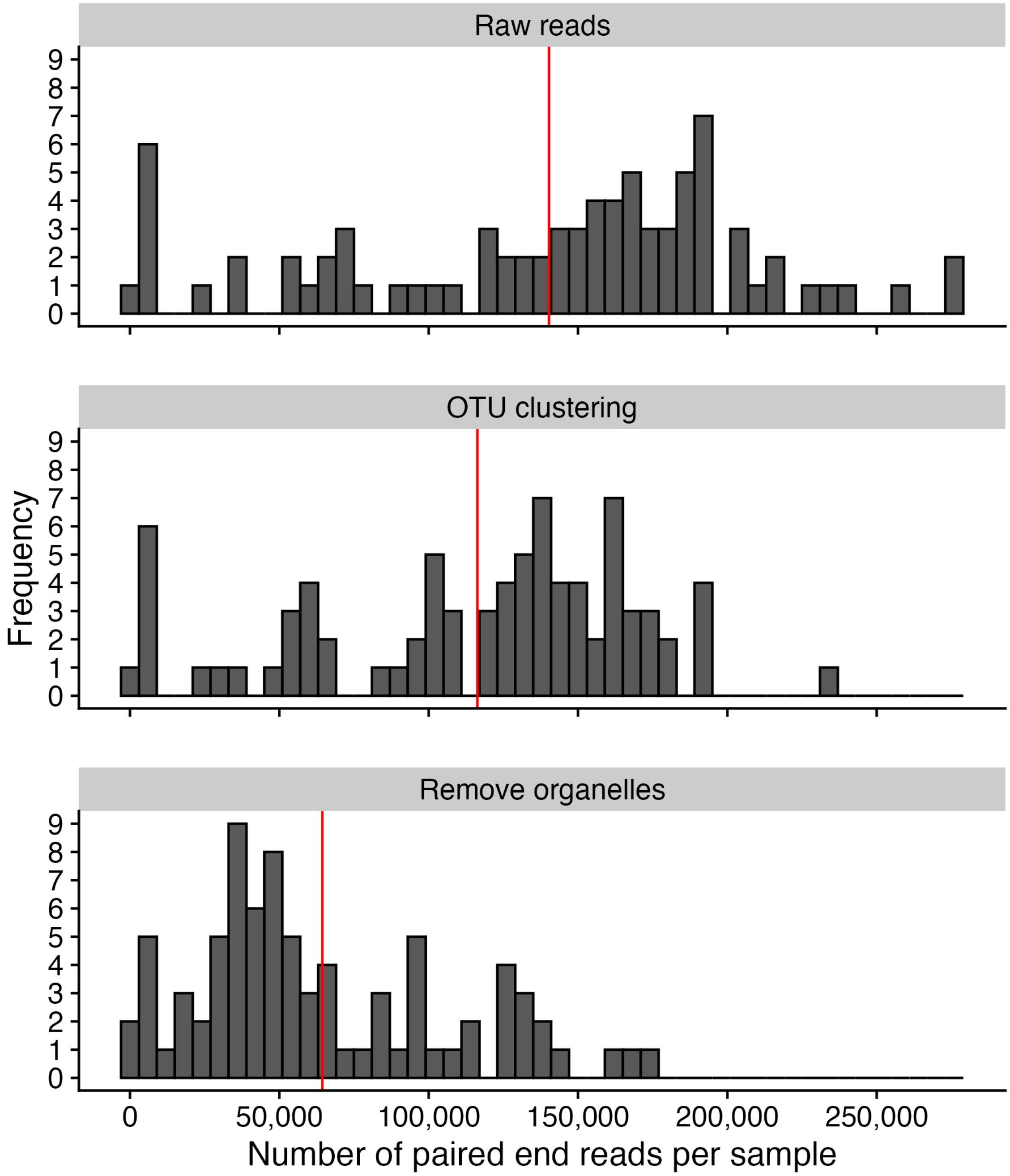


Supplemental Figure 1. Distribution of reads per sample. The vertical red line represents the mean in all panels. Top panel, distribution of raw read count per sample. Middle panel, distribution of read count per sample after OTU clustering (i.e., reads that correspond to OTUs after generating a feature table with QIIME2). Bottom panel, distribution of read count per sample after mitochondrion- and chloroplast-aligning reads were removed from the analysis.

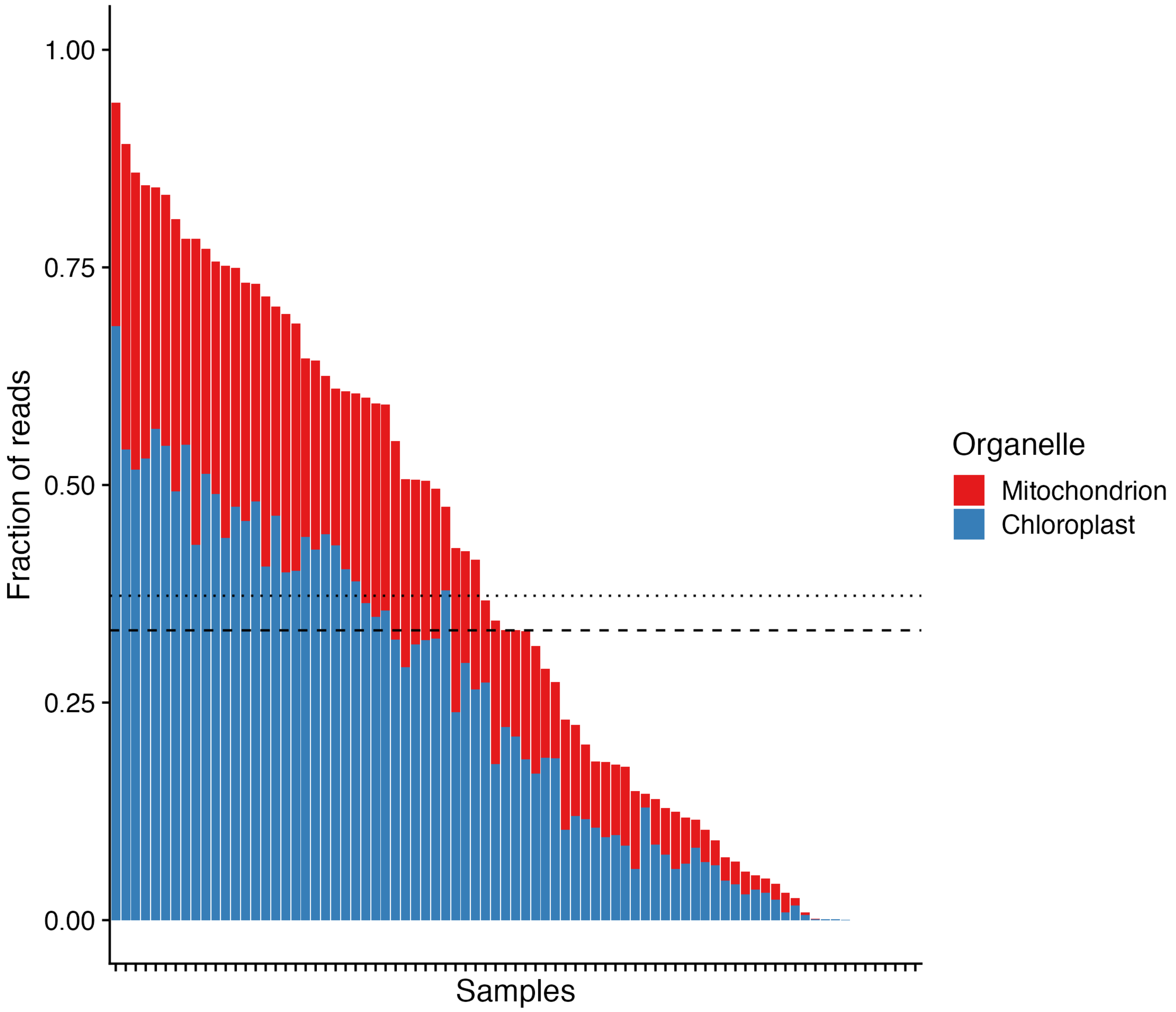

Supplemental Figure 2. Fraction of reads per sample aligning to organellar DNA. Dashed line, median fraction; Dotted line, mean fraction.

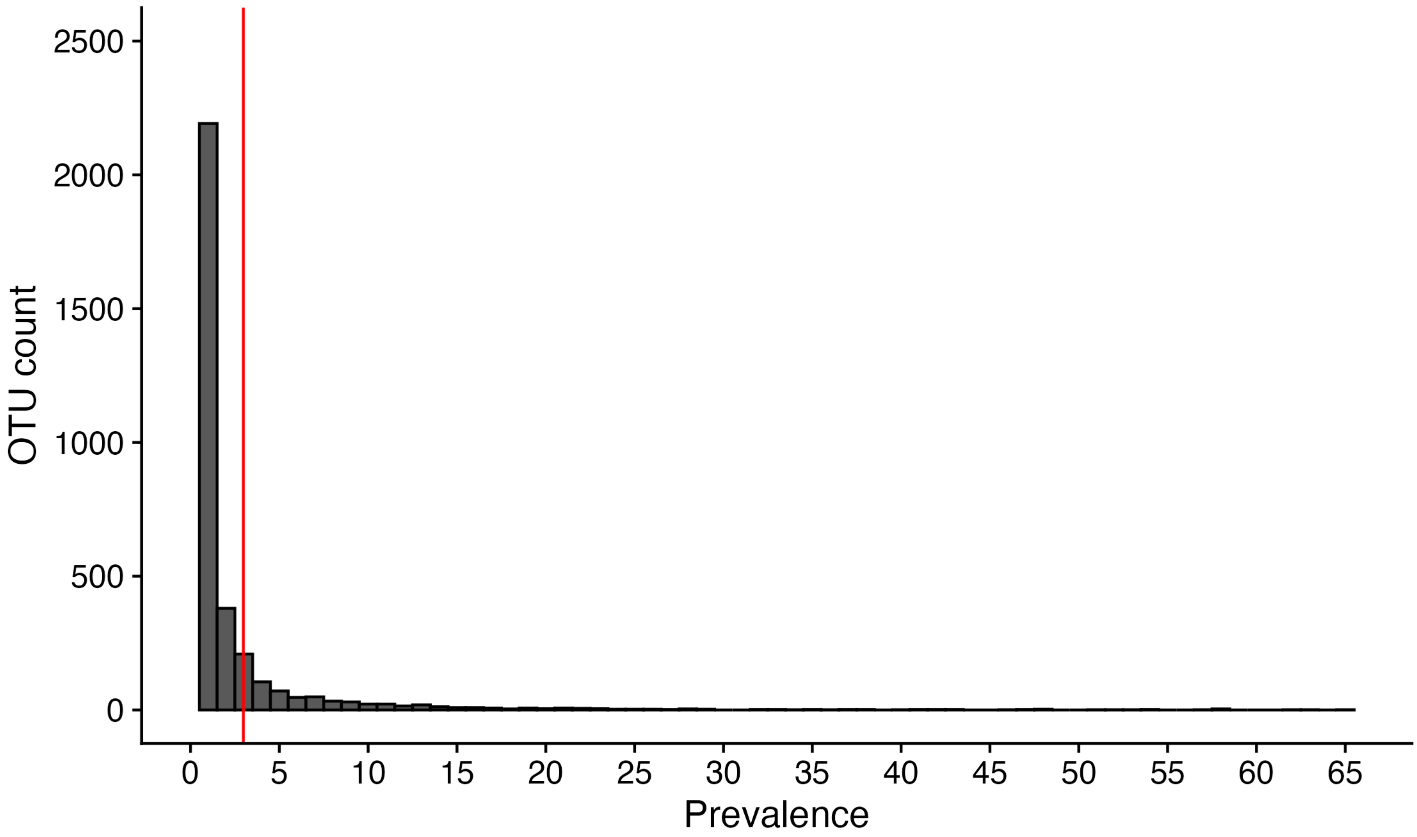


Supplemental Figure 3. Distribution of OTU prevalence. The red vertical line represents the mean. No OTU was present in all samples.

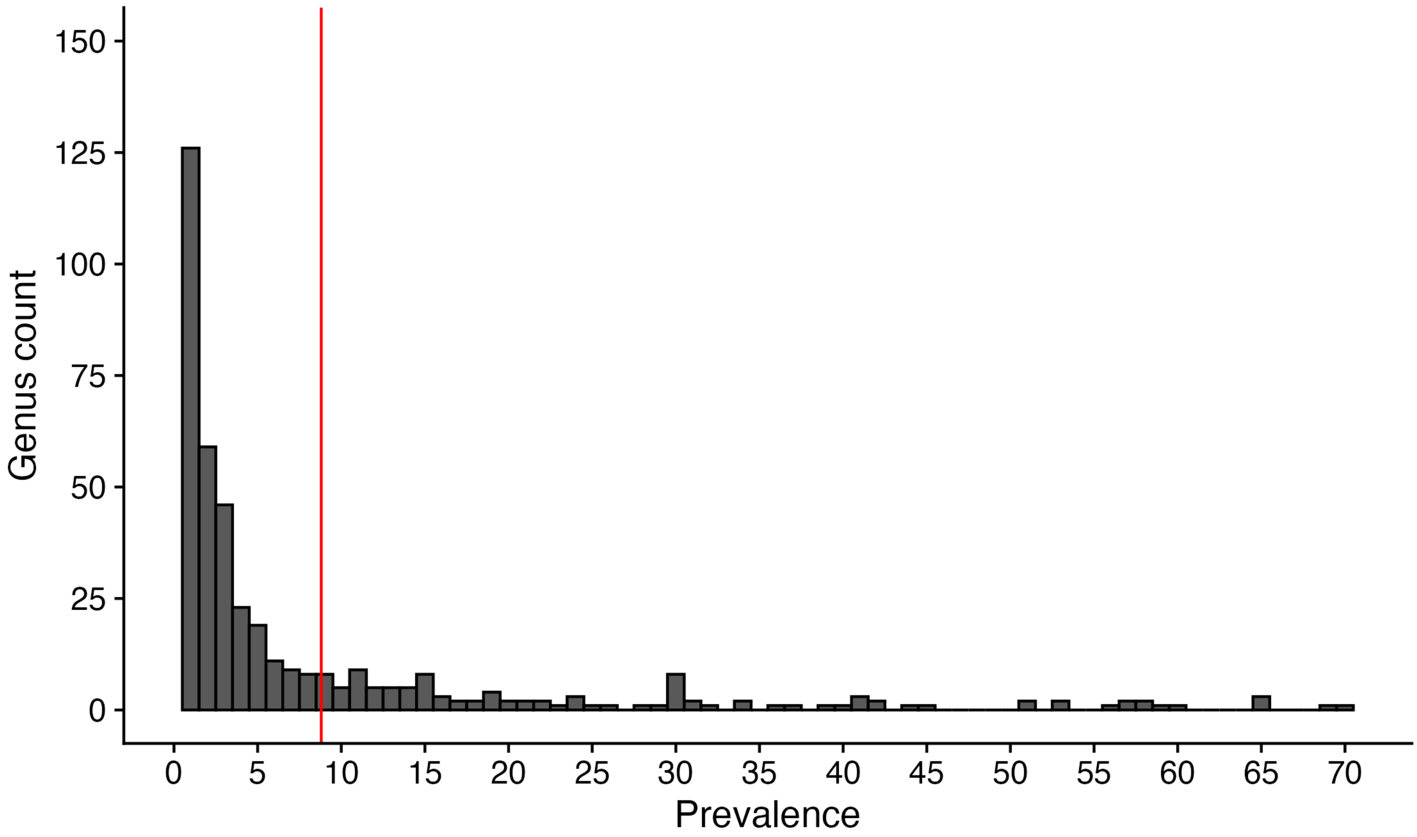


Supplemental Figure 4. Distribution of genus prevalence. The red vertical line represents the mean.

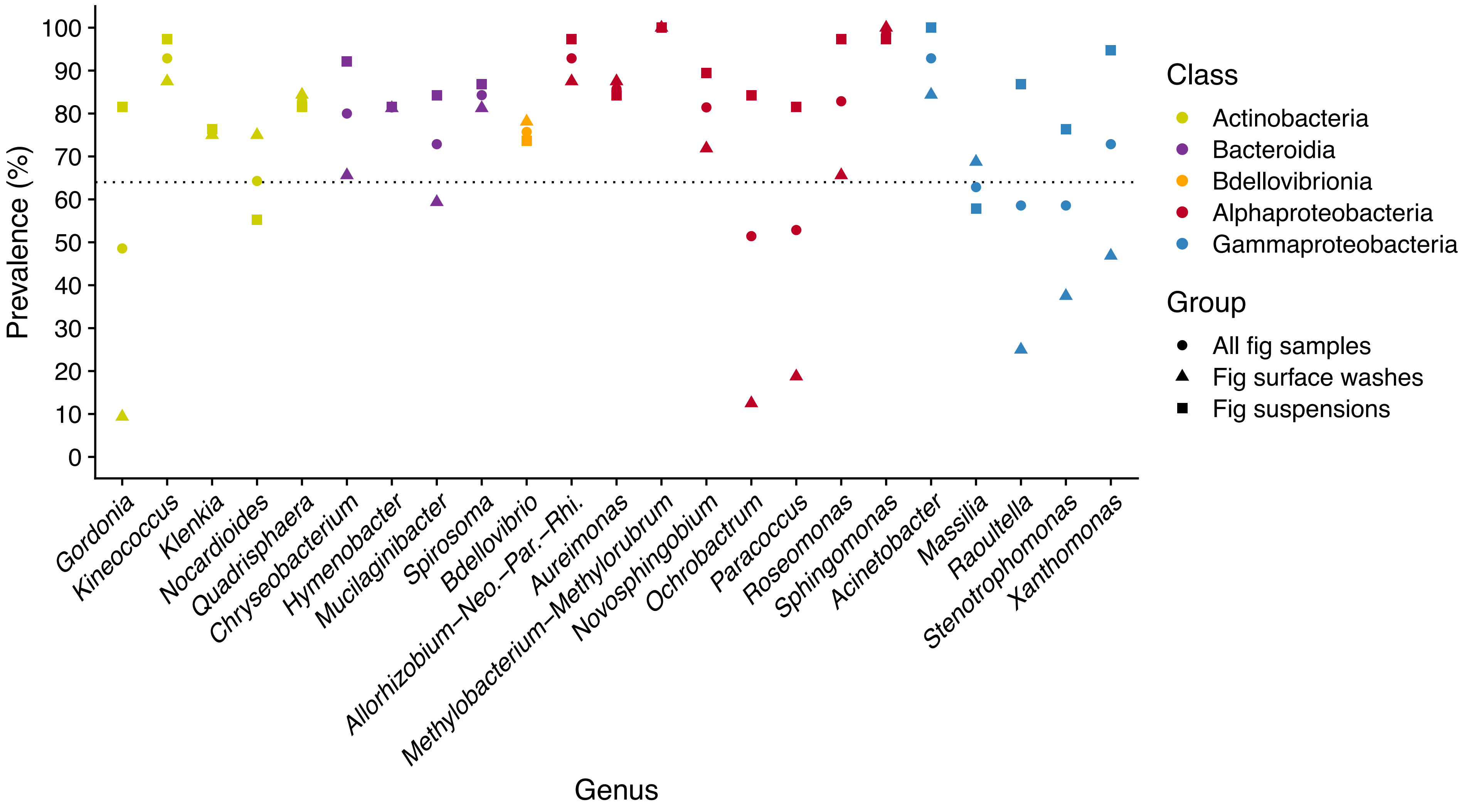


Supplemental Figure 5. Genera with >65% prevalence in a given group (All fig samples, fig surface washes, or fig suspensions).

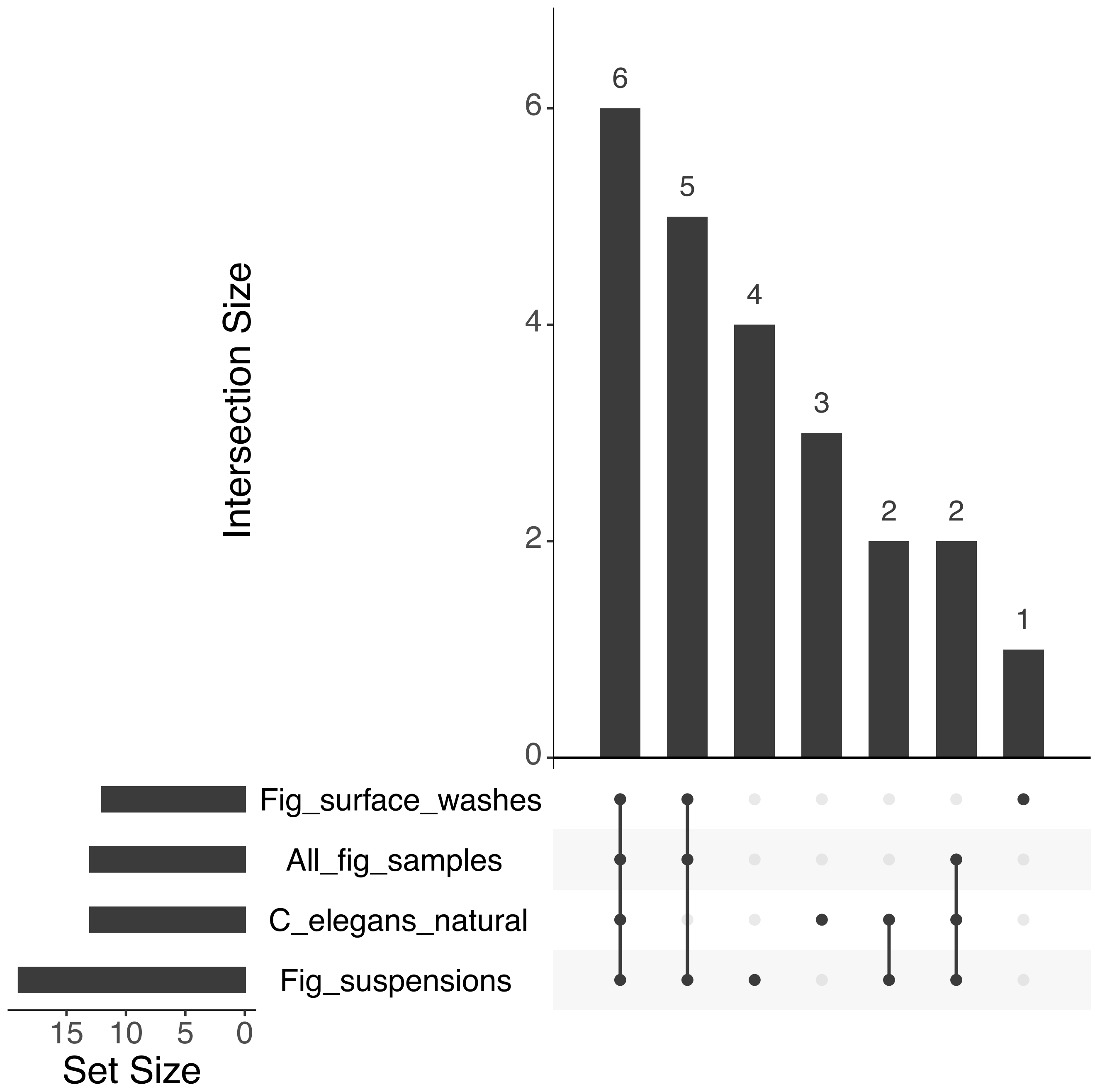


Supplemental Figure 6. Upset plot of prevalent families among fig samples and the most prevalent families in *C. elegans* microbial communities as reported in Zhang et al. 2017. *Frontiers in Microbiology*. Families prevalent among both fig samples and *C. elegans* samples: *Microbacteriaceae*, *Comamonadaceae*, *Moraxellaceae*, *Xanthomonadaceae*, *Acetobacteraceae*, *Sphingomonadaceae*, *Rhodobacteraceae*, *Sphingobacteriaceae*, *Weeksellaceae*, and *Enterobacteriaceae*.

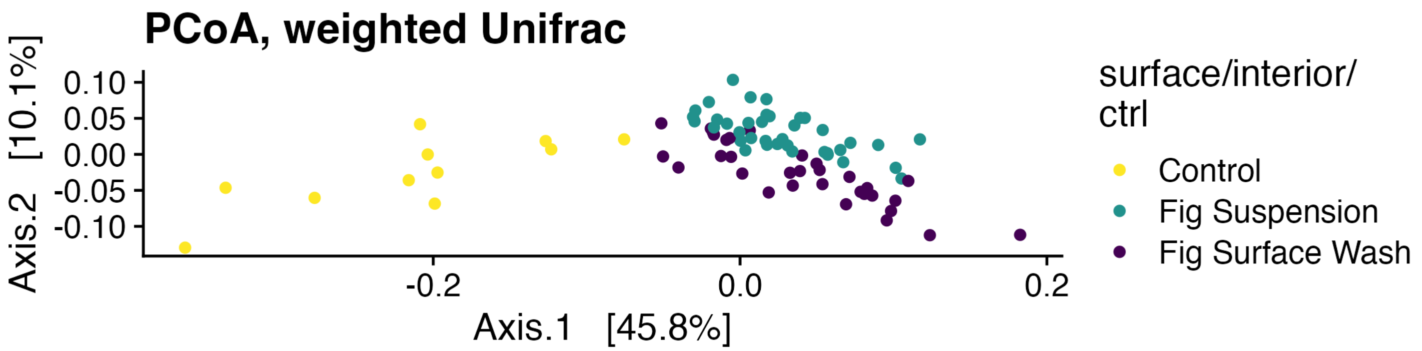


Supplemental Figure 7. Principal coordinates analysis (PCoA), weighted UniFrac distances, including controls.

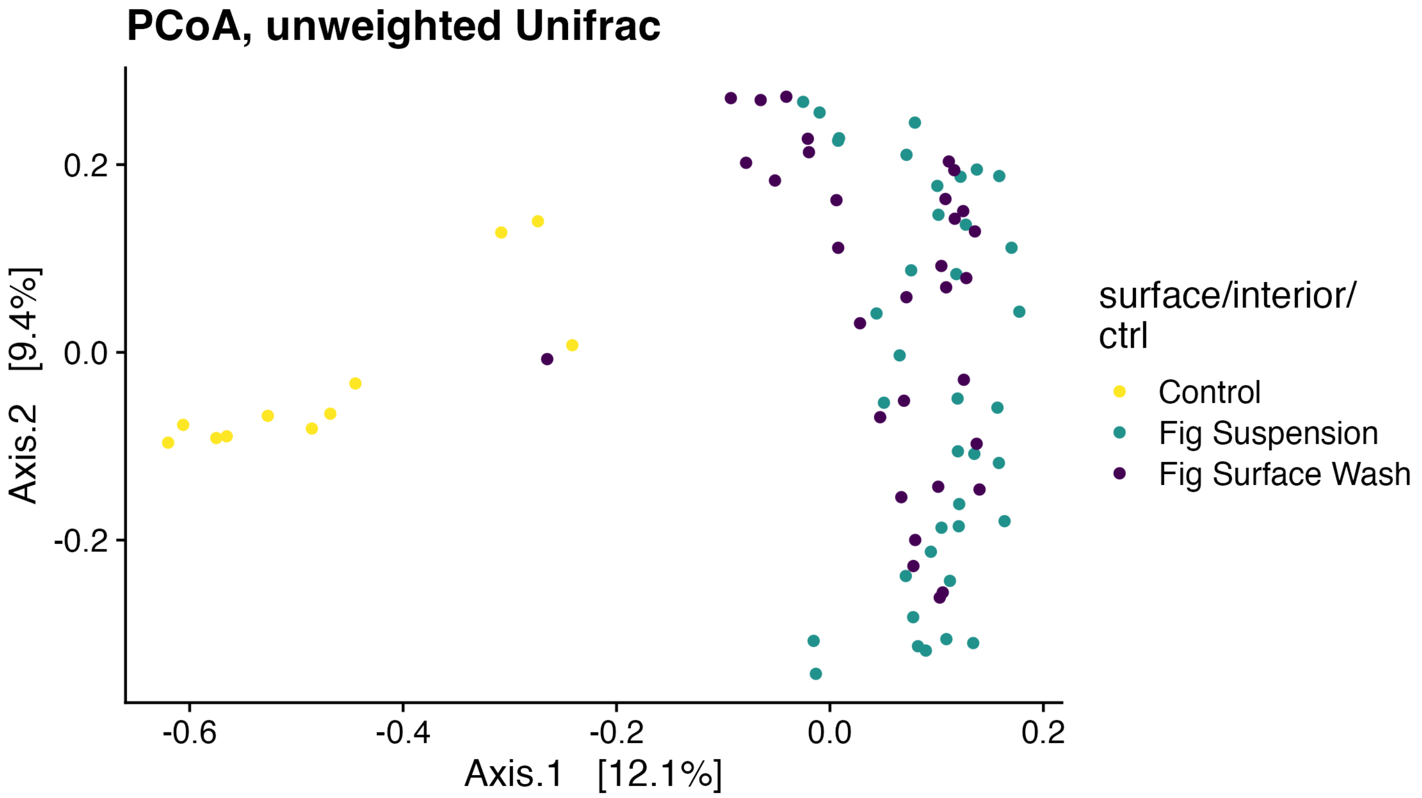


Supplemental Figure 8. Principal coordinates analysis (PCoA), unweighted UniFrac distances, including controls.

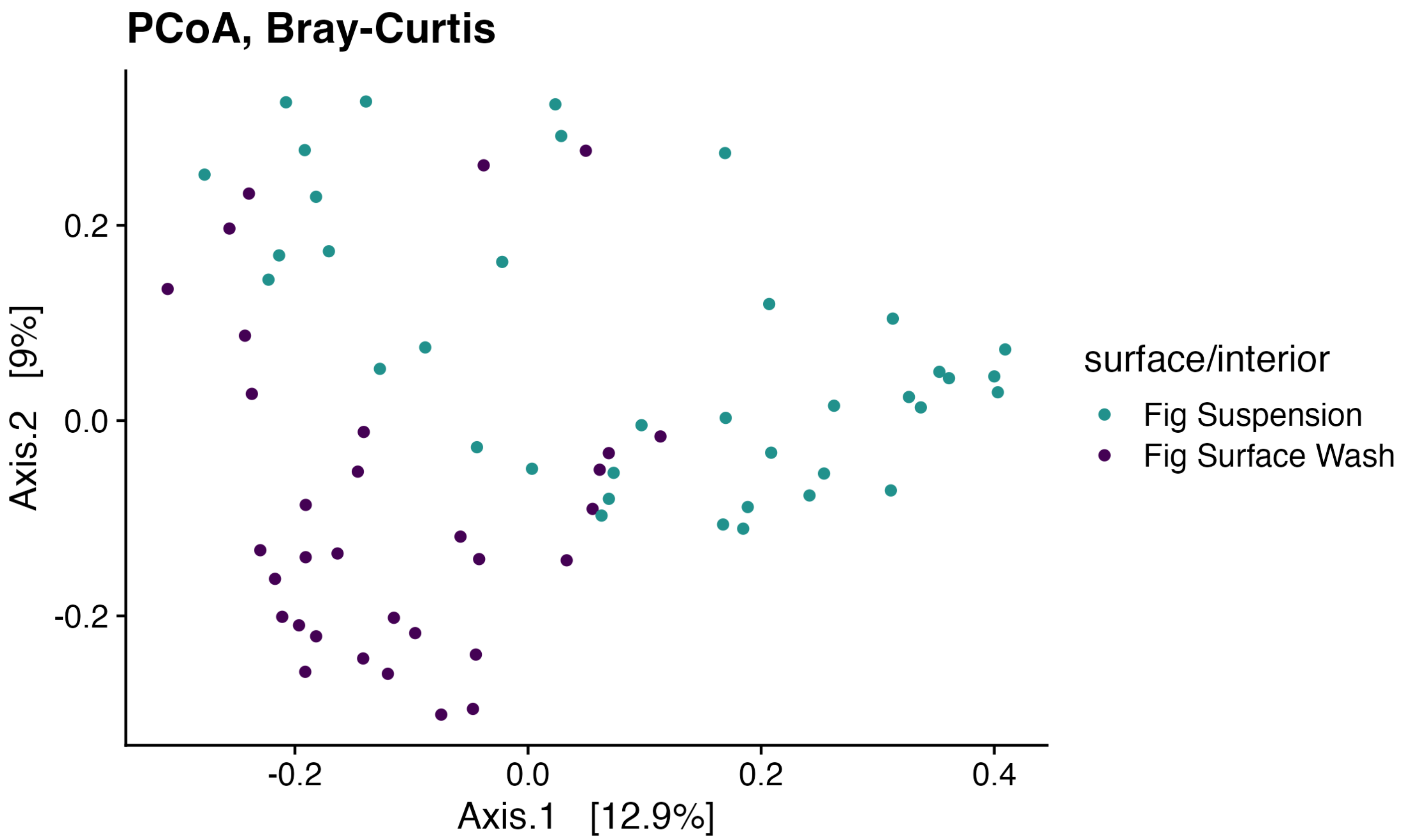


Supplemental Figure 9. Principal coordinates analysis (PCoA), Bray-Curtis distances, not including controls.

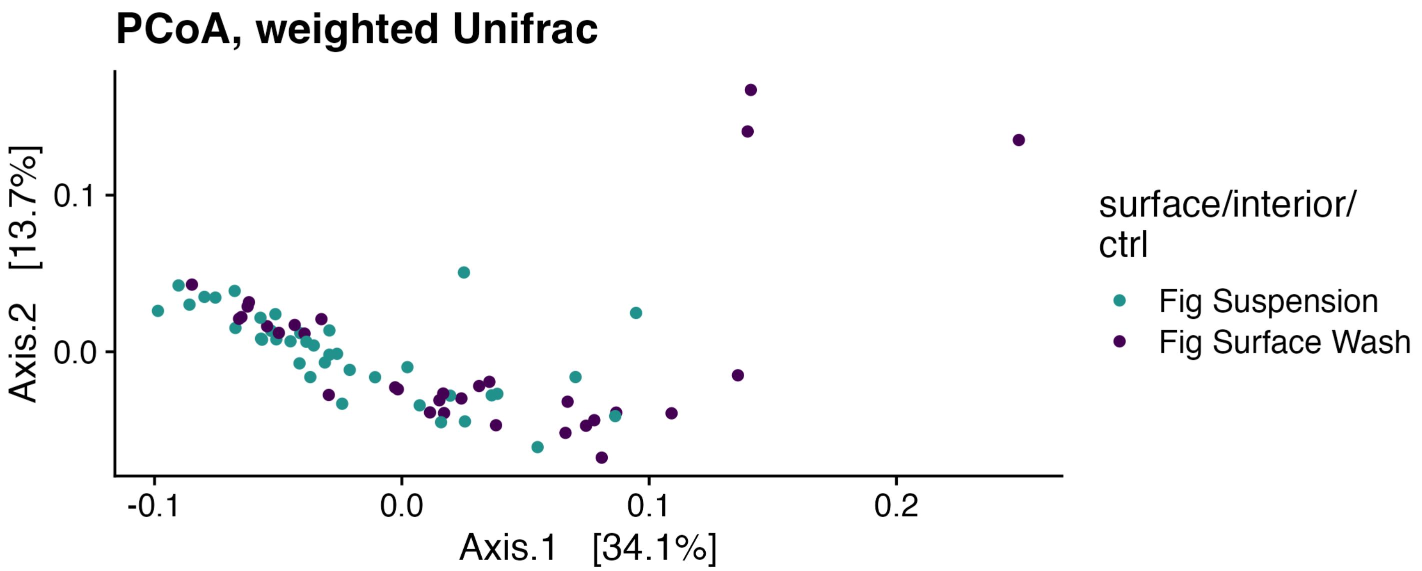


Supplemental Figure 10. Principal coordinates analysis (PCoA), weighted Unifrac distances, not including controls.

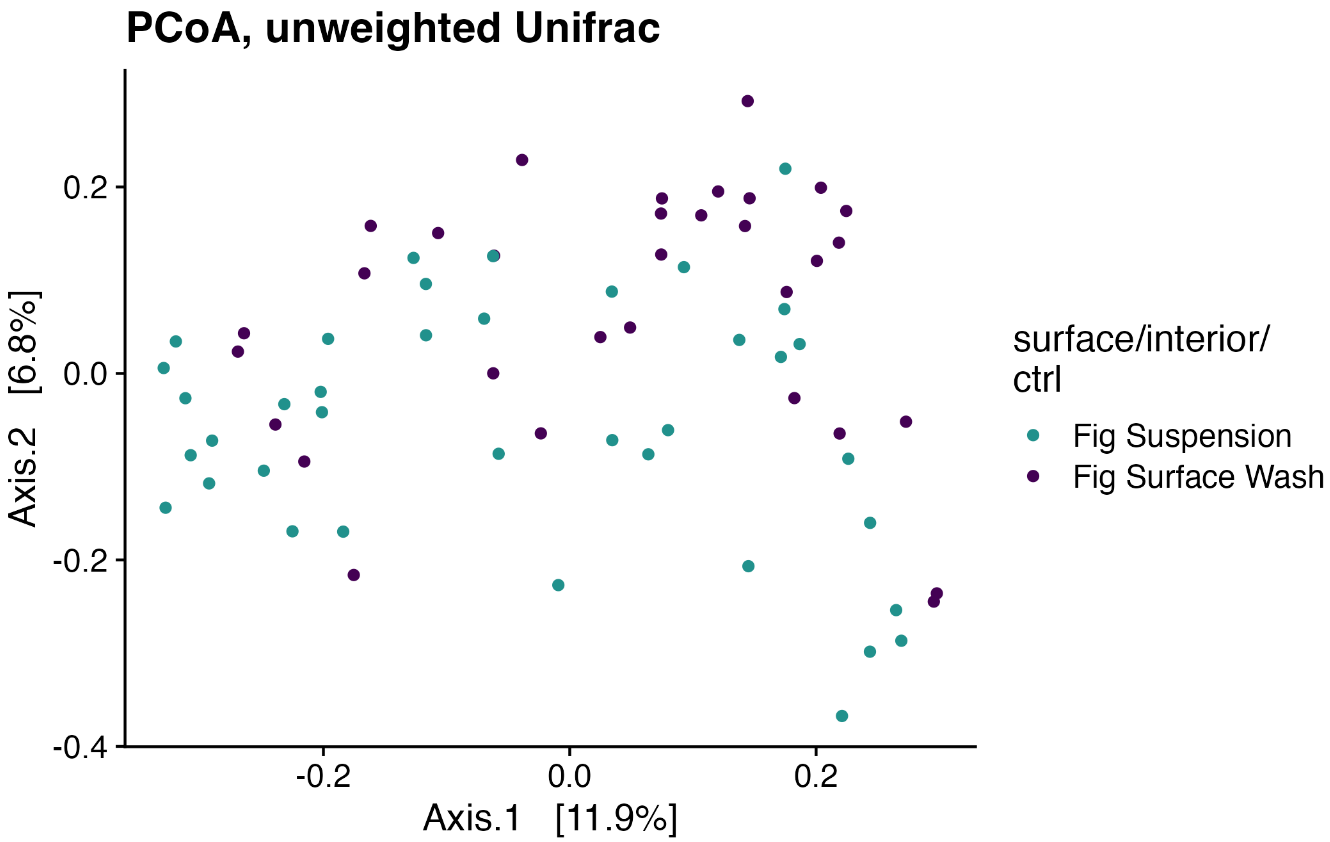


Supplemental Figure 11. Principal coordinates analysis (PCoA), unweighted Unifrac distances, not including controls.

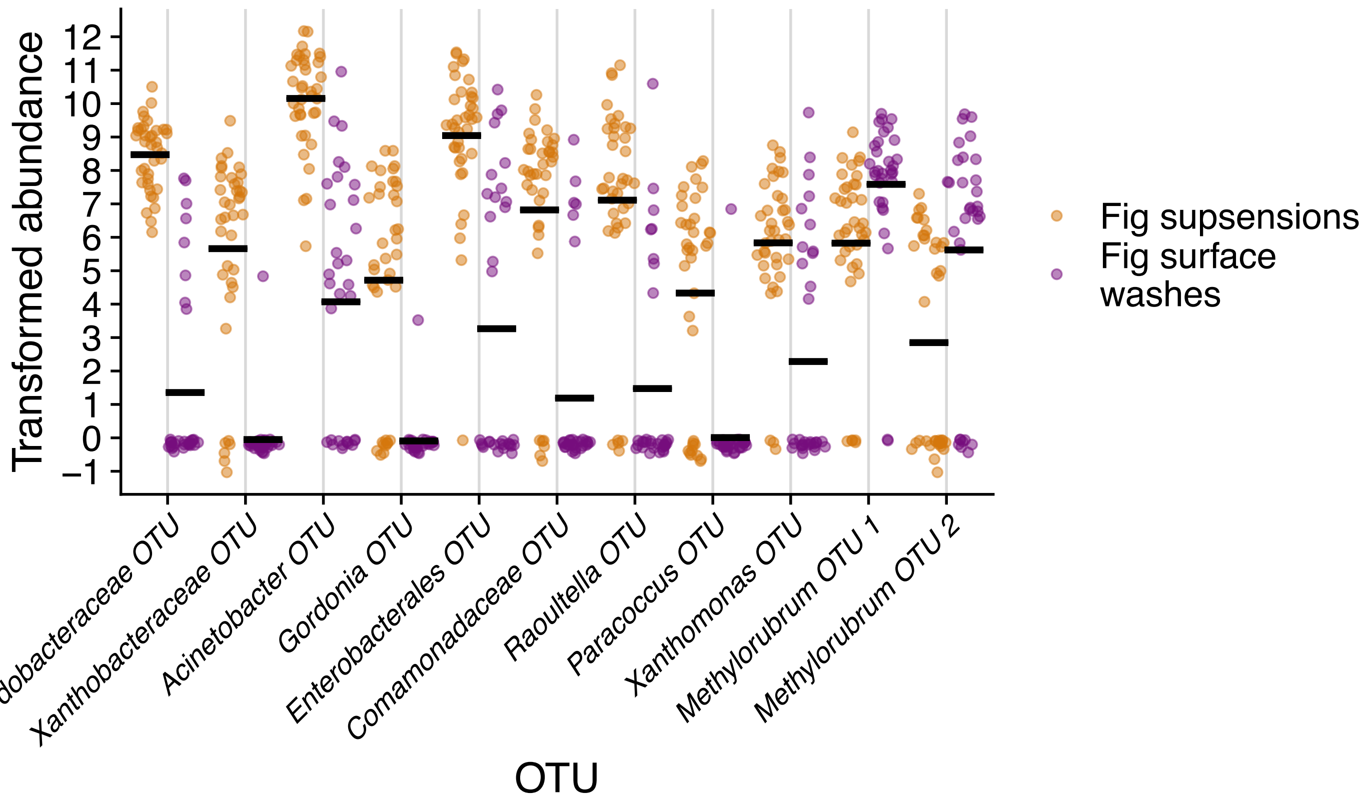


Supplemental Figure 12. OTUs differentially abundant among fig suspensions and fig surface washes. Plotted are centered log-ratio transformed abundances by sample type (fig suspensions or surface washes). The eleven significant OTU are plotted and ordered by effect size. Sina plots are strip charts that take the contours of a violin plot. Horizontal bars represent means.

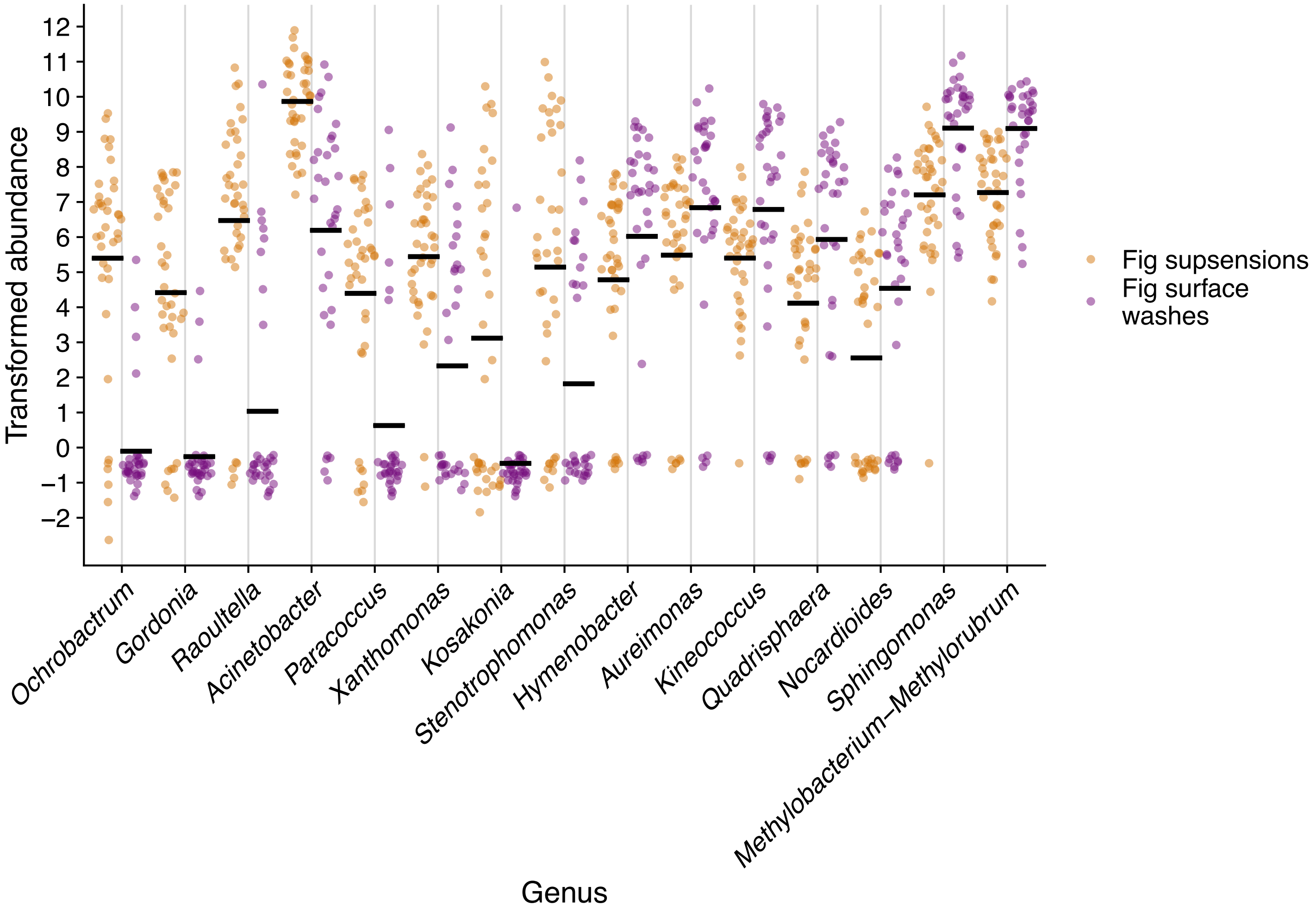


Supplemental Figure 13. Genera differentially abundant among fig suspensions and fig surface washes. Plotted are centered log-ratio transformed abundances by sample type (fig suspensions or surface washes). The fifteen significant genera (not including one labeled "Uncultured") are plotted and ordered by effect size. Sina plots are strip charts that take the contours of a violin plot. Horizontal bars represent means.

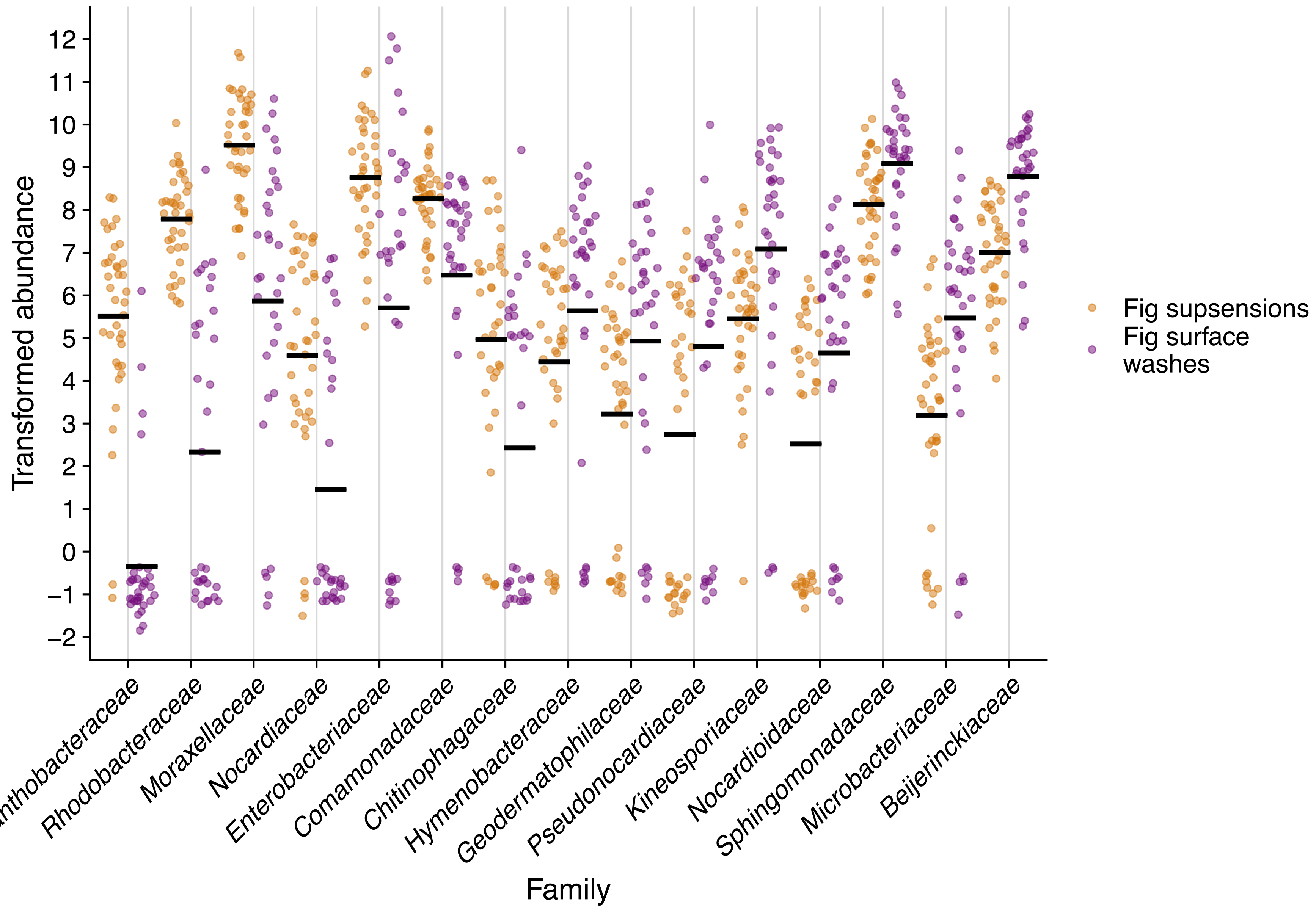


Supplemental Figure 14. Families differentially abundant among fig suspensions and fig surface washes. Plotted are centered log-ratio transformed abundances by sample type (fig suspensions or surface washes). The fifteen significant families (not including one labeled "Uncultured") are plotted and ordered by effect size. Sina plots are strip charts that take the contours of a violin plot. Horizontal bars represent means.

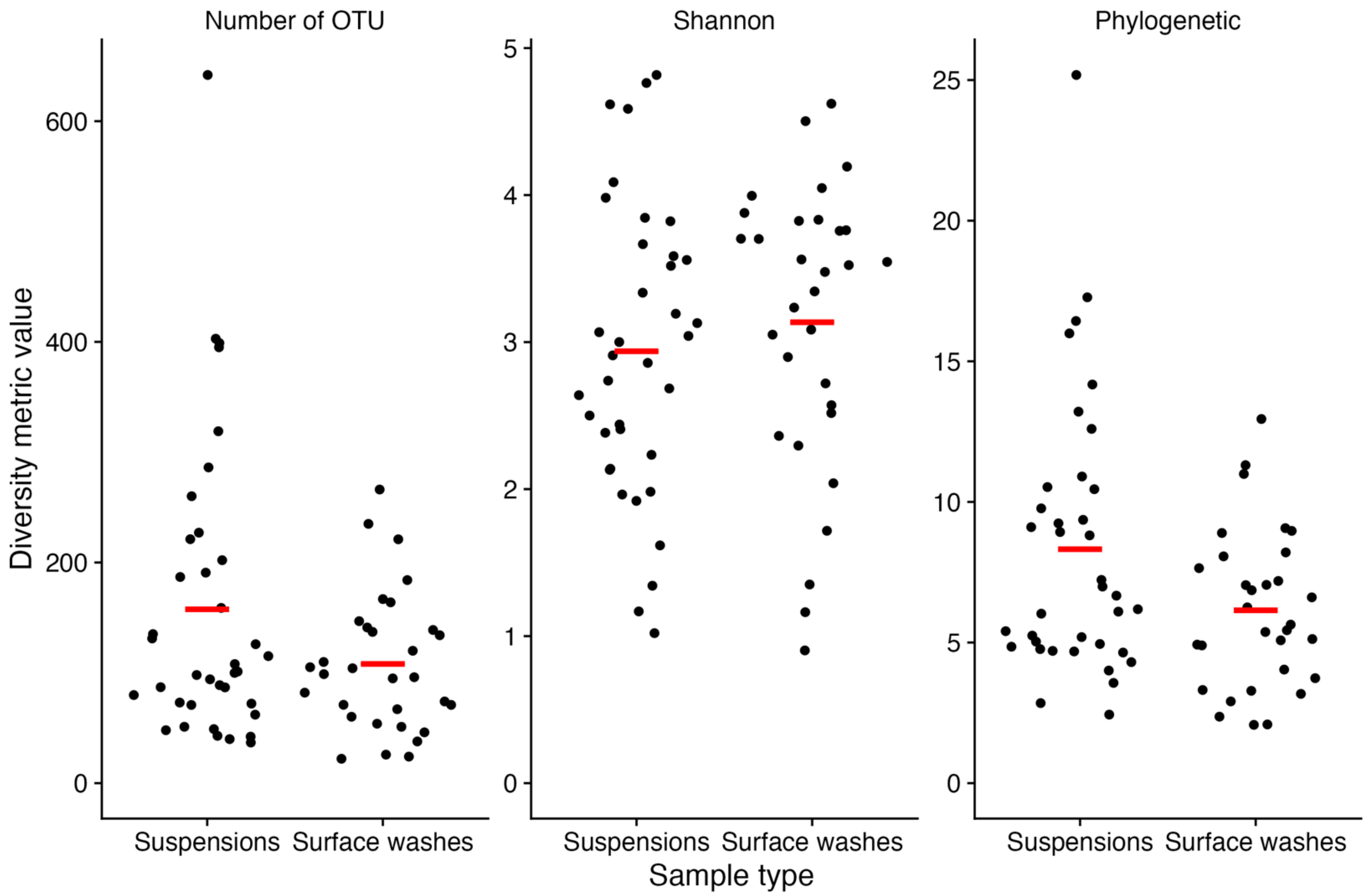


Supplemental Figure 15. Microbial α-diversity is comparable among fig suspensions and fig surface washes. Three diversity metrics (number of OTU's found in each sample; Shannon diversity; and Phylogenetic diversity) are plotted. Sina plots are strip charts with points taking the contours of a violin plot. Horizontal bars represent means.

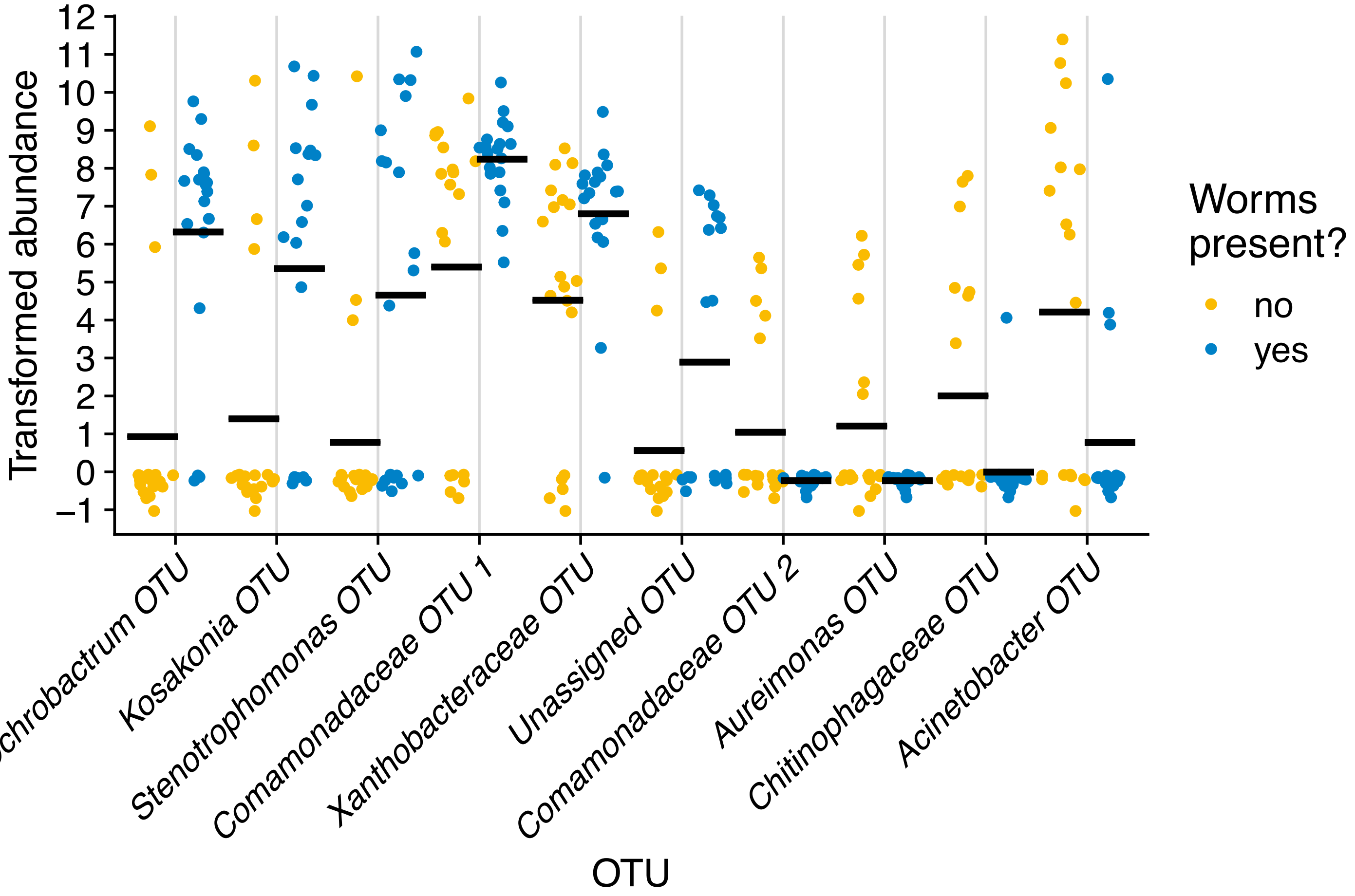


Supplemental Figure 16. OTUs differentially abundant among fig suspensions with and without nematodes. Plotted are centered log-ratio transformed abundances by sample type (fig suspensions or surface washes). The ten OTU with the lowest unadjusted Mann-Whitney U *p*-values are plotted and sorted by effect size (For the *Ochobactrum* OTU, the FDR adjusted *p*=0.21; for all others, the FDR adjusted *p*=0.91). Sina plots are strip charts that take the contours of a violin plot. Horizontal bars represent means.

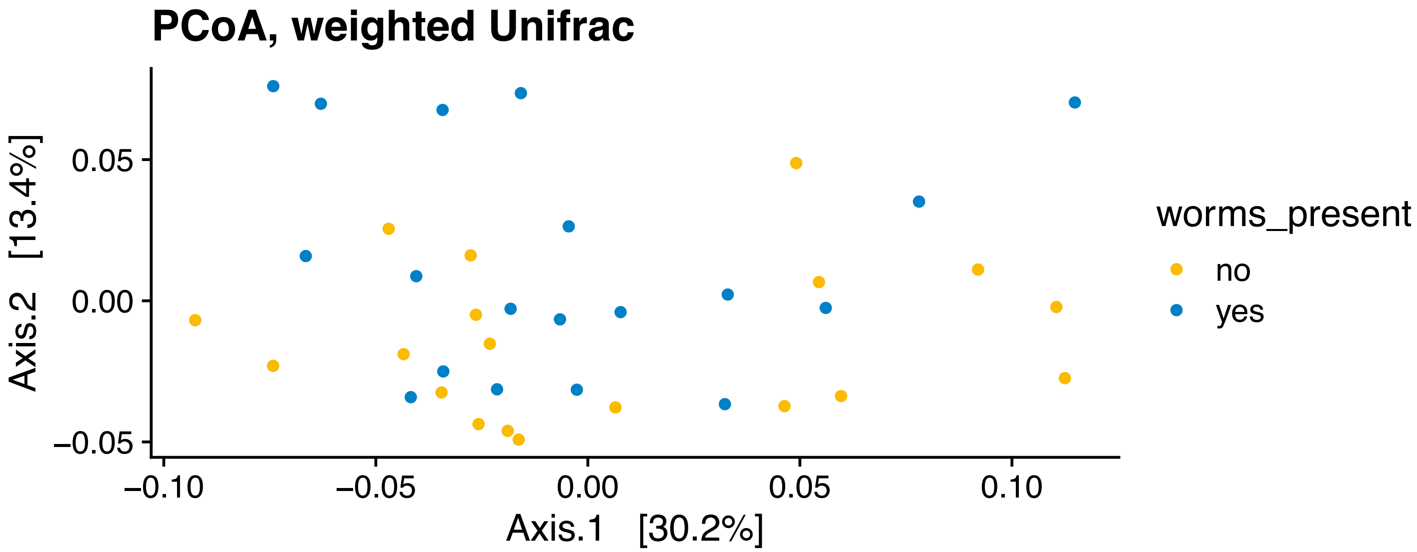


Supplemental Figure 17. Principal coordinates analysis (PCoA), weighted Unifrac distances, only including fig suspensions.

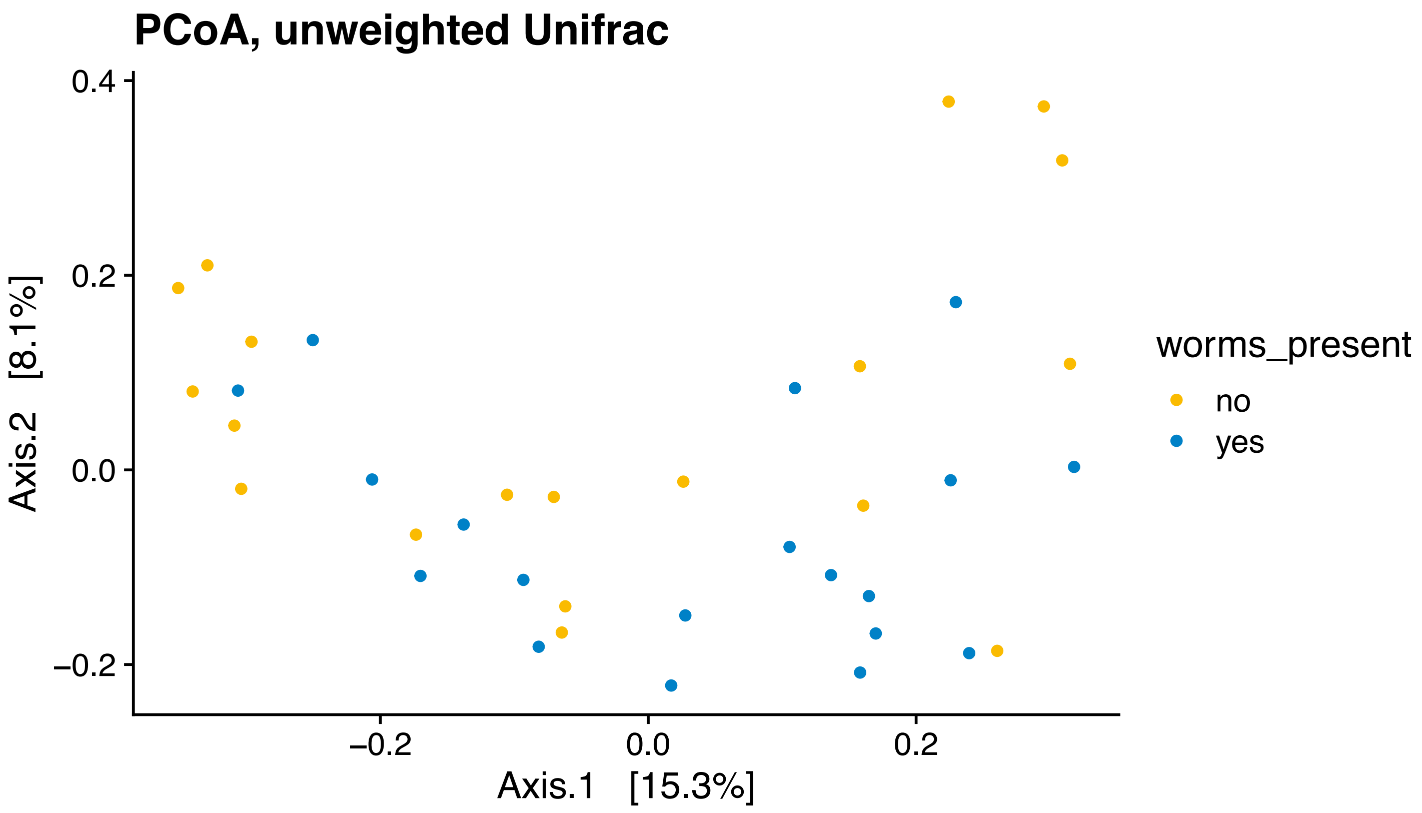


Supplemental Figure 18. Principal coordinates analysis (PCoA), unweighted Unifrac distances, only including fig suspensions.

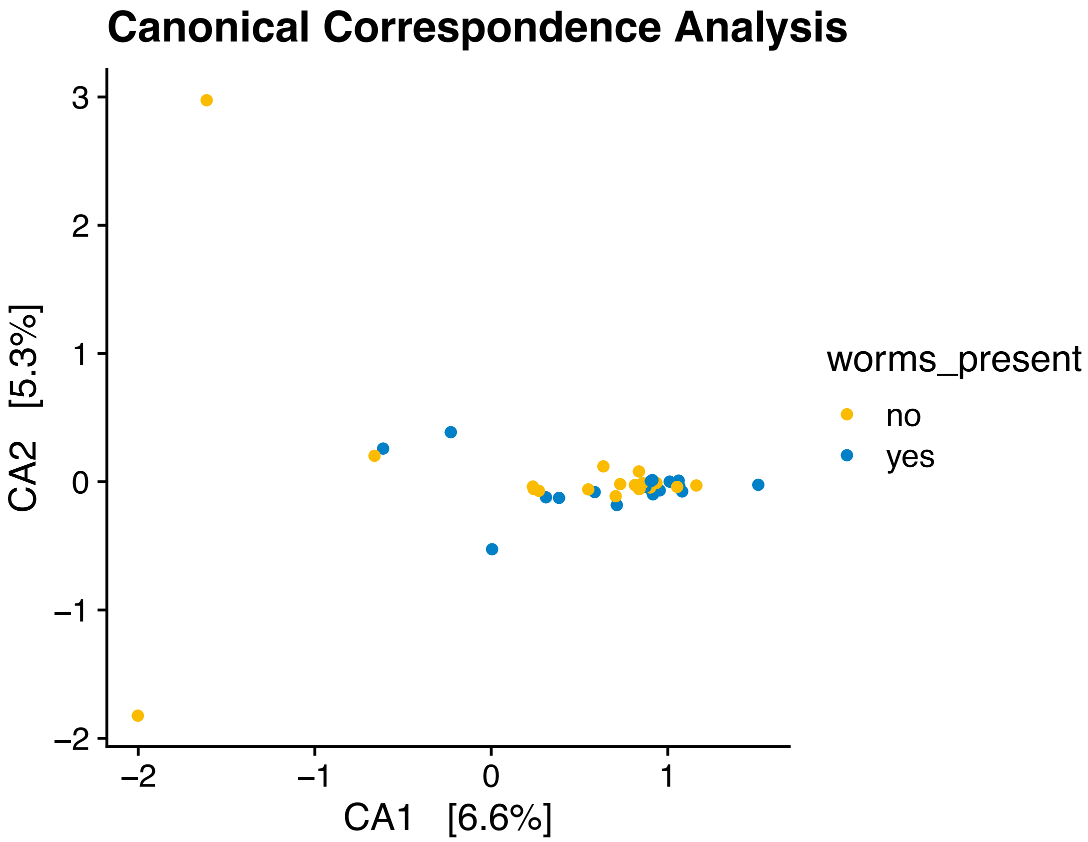


Supplemental Figure 19. Canonical correspondence analysis, only including fig suspensions.

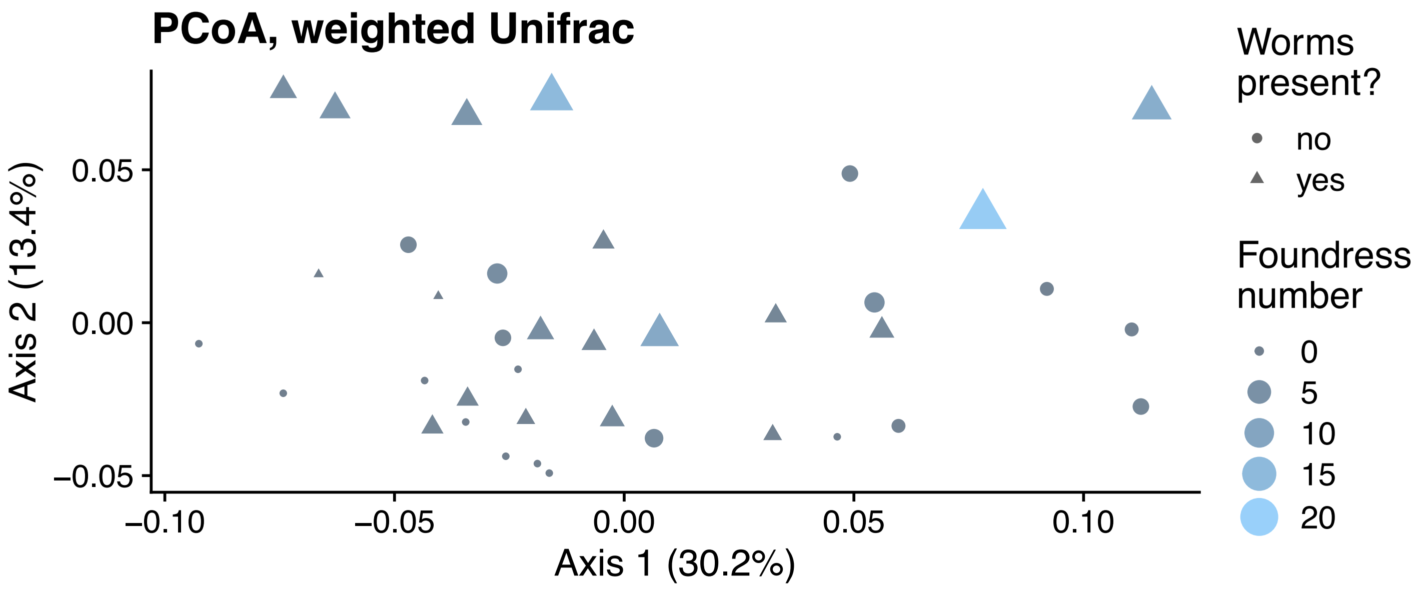


Supplemental Figure 20. Principal coordinates analysis (PCoA), weighted Unifrac distances, only including fig suspensions. Here, with points sized by foundress wasp number.

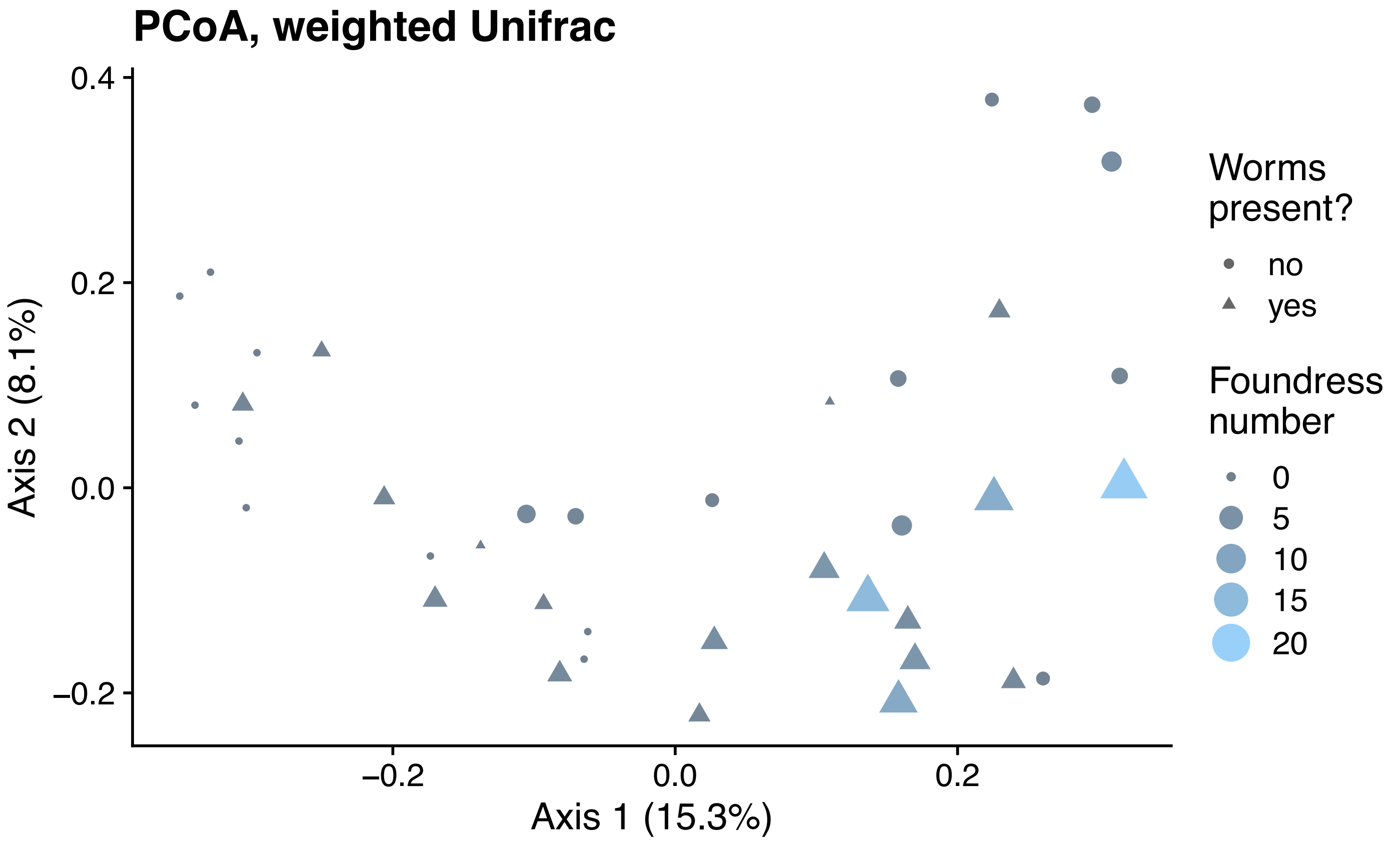


Supplemental Figure 21. Principal coordinates analysis (PCoA), weighted Unifrac distances, only including fig suspensions. Here, with points sized by foundress wasp number.

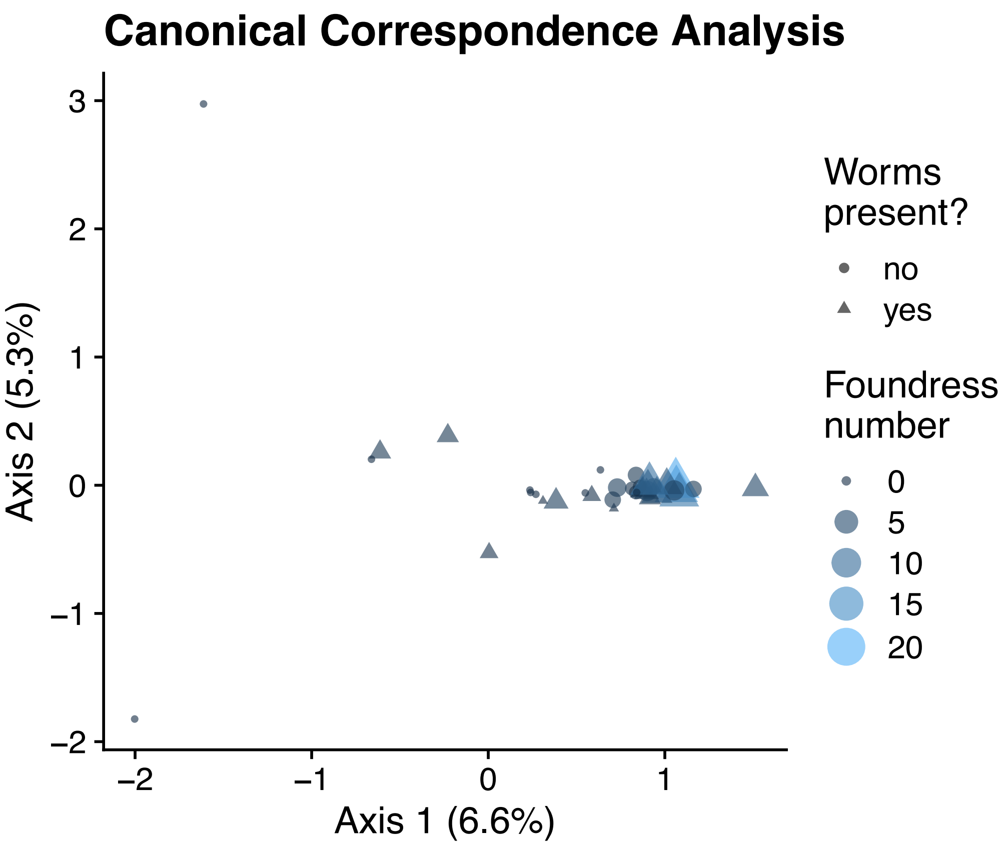


Supplemental Figure 22. Canonical correspondence analysis, only including fig suspensions. Here, with points sized by foundress wasp number.

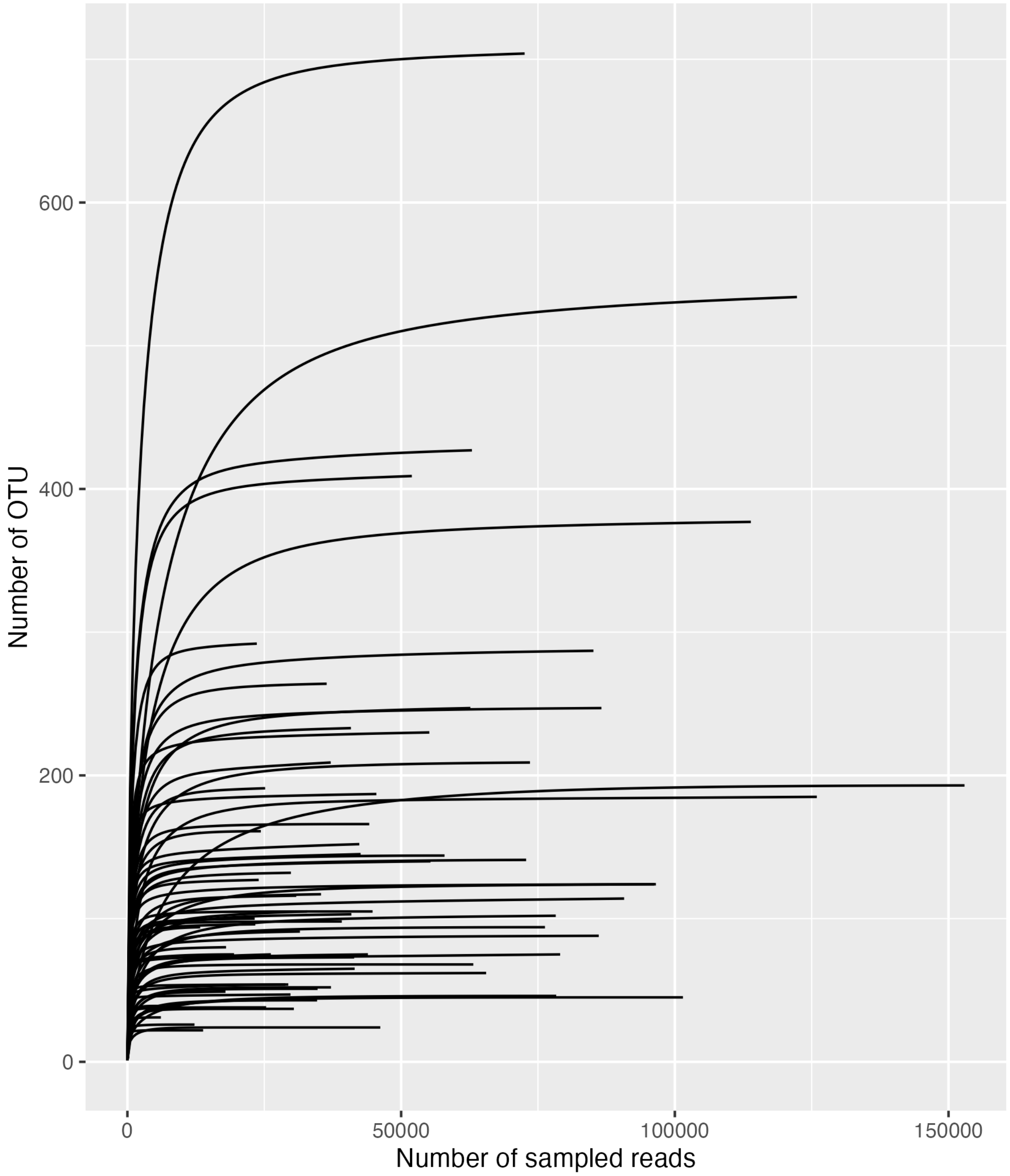


Supplemental Figure 23. Rarefaction curves. Here, showing the entire distribution of read counts.

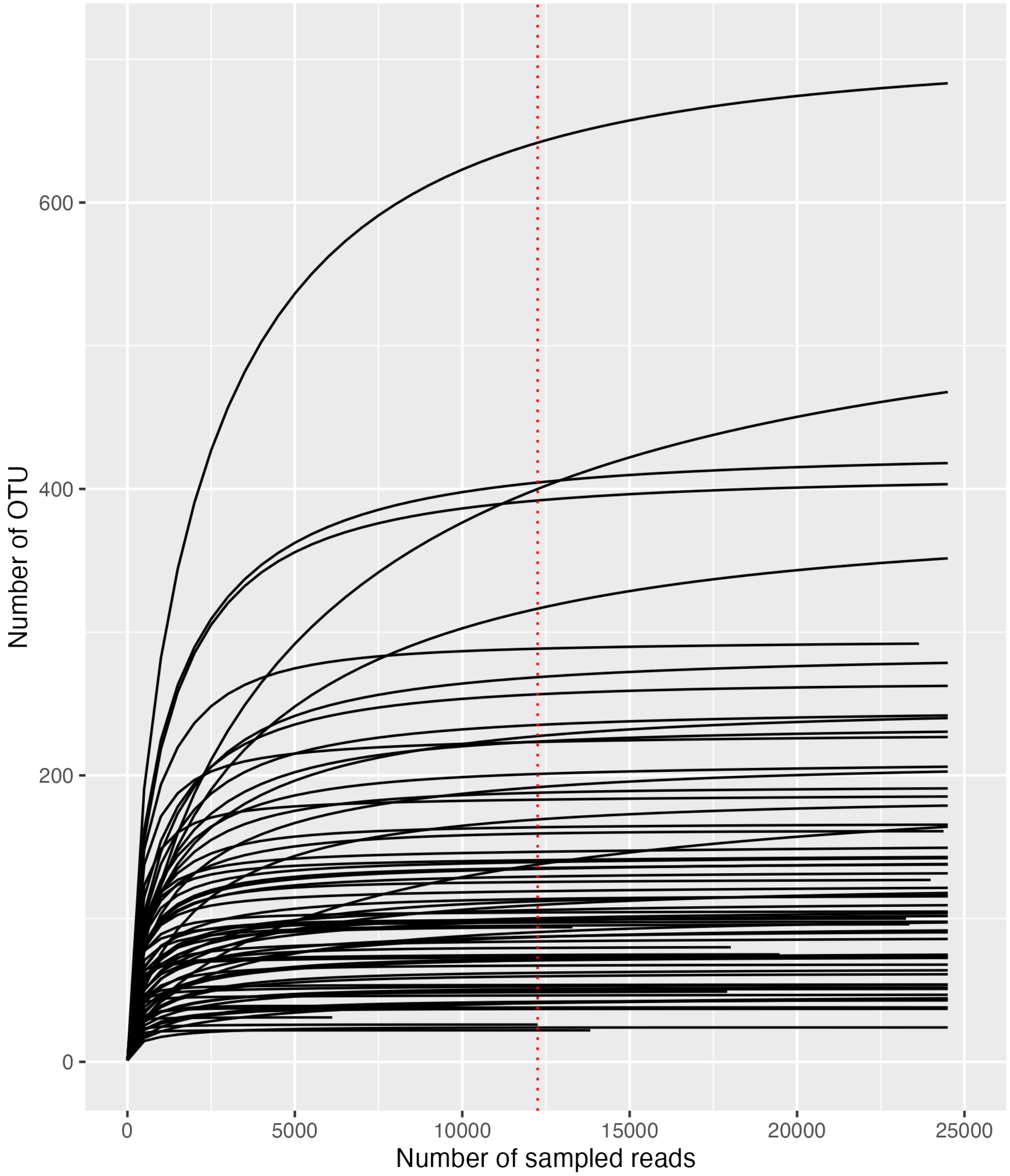


Supplemental Figure 24. Rarefaction curves. Here, the x-axis is bounded at 25,000 reads. The vertical dotted line denotes the read count used for rarefaction in estimates of α-diversity (12,252).

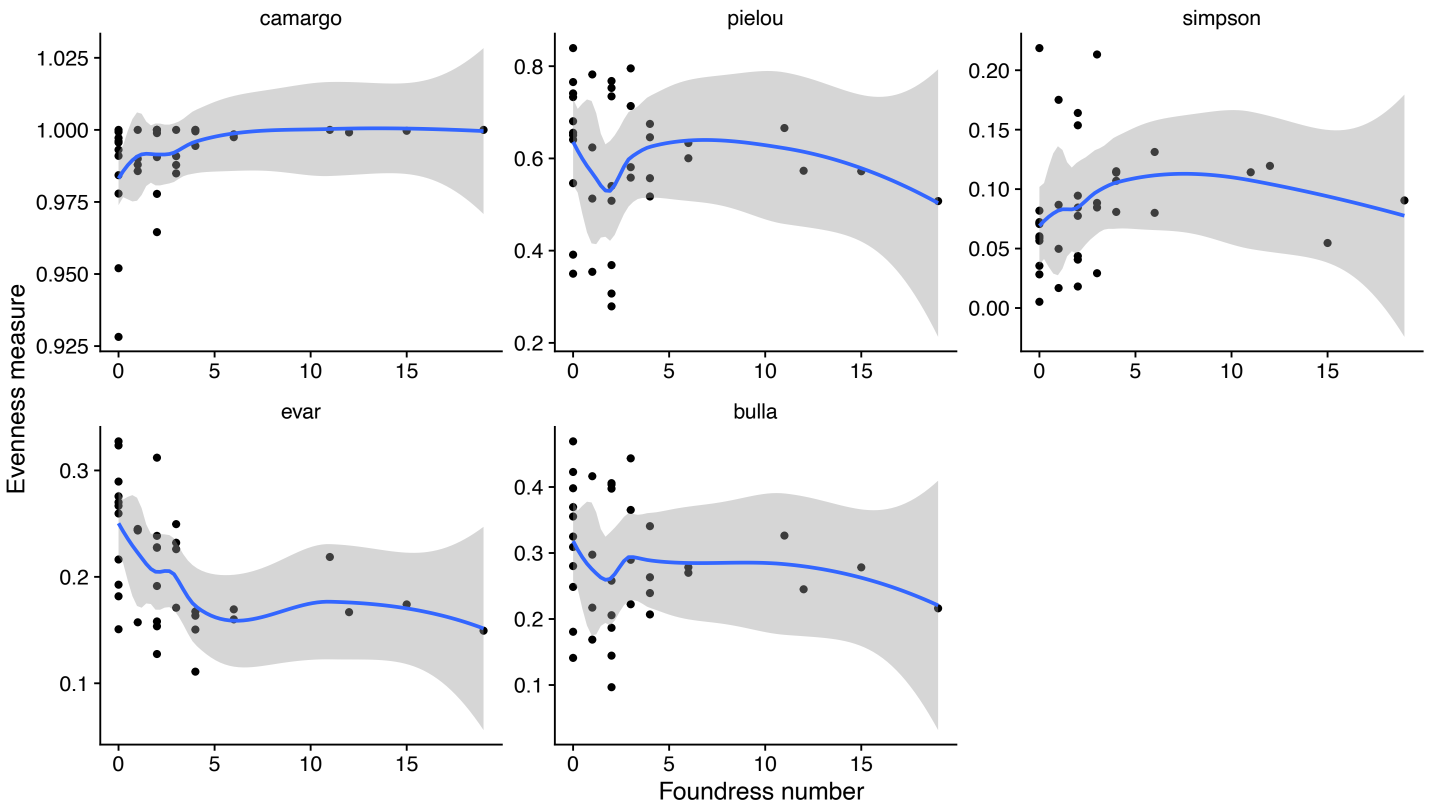


Supplemental Figure 25. The relationship between microbial community evenness and foundress number. "carmago": Camargo's evenness; "pielou": Pielou's evenness or the Shannon or Shannon-Weaver/Wiener/Weiner evenness, H/ln(S); "simpson": Simpson’s evenness (inverse Simpson diversity/S); "evar": Smith and Wilson’s Evar index; "bulla": Bulla’s index (O). Only Smith and Wilson’s Evar index reveals a significant relationship between evenness and foundress number (OLS p=0.012); all others, p=0.066-0.64).
